# Dating the photosynthetic organelle evolution in Archaeplastida, *Paulinella* and secondary-plastid bearing lineages

**DOI:** 10.1101/2022.07.01.497312

**Authors:** Filip Pietluch, Paweł Mackiewicz, Katarzyna Sidorczuk, Przemysław Gagat

## Abstract

Photosynthetic eukaryotes have shaped the Earth’s biosphere by producing O_2_ and converting light into organic compounds in specialized organelles called plastids. Plastids evolved from free-living cyanobacteria engulfed by heterotrophic unicellular eukaryotes in processes called cyanobacterial endosymbioses. Two independent such processes have been reported so far. The first gave rise to primary plastids and three Archaeplastida lineages: glaucophytes, red algae and green algae with land plants, whereas the second resulted in chromatophores in the rhizarian amoeba *Paulinella*. Importantly, archaeplastidans donated their plastids to many protist groups, thereby further spreading photosynthesis across the tree of life. To reveal the complex plastid evolution, we performed comprehensive phylogenetic and multi-clock analyses based on new fossil calibration points and the greatest number yet of plastid-encoded proteins from 108 taxa, representing a large diversity of photosynthetic organisms. Our results indicate that primary plastids evolved prior to 2.1 - 1.8 Bya, i.e. before glaucophytes diverged from the other archaeplastidans. Like the primary plastids before, *Paulinella* chromatophores evolved in low salinity habitats and possibly before 292 - 266 Mya. Red and green algae were engulfed by cryptophyte and chlorarachniophyte ancestors between 1.7 - 1.4 Bya, and 1.1 - 1.0 Bya, respectively; the former subsequently triggered plastid transfers to other eukaryotes. The diversification rate of the photosynthetic organisms increased with temperature and CO_2_ but decreased with O_2_ and volcanic activity. We also studied the impact of various molecular clocks and calibration sets on the age estimation and clearly indicate that the clocks are the source of greater differences.

**Significance Statement:** Cyanobacteria and eukaryote endosymbioses created a multitude of photosynthetic organelles called plastids that feed most life on our planet. For decades scientists have been trying to untangle the puzzle of plastid evolution, i.e. when and how plastids were acquired and spread throughout the eukaryotic tree of life. To answer these questions we applied phylogenetic and multi-clock methods combined with new fossil calibration points on large data sets. Our results push back in the Earth’s history most key events concerning plastid evolution compared to previous reports. Additionally, we discovered a significant impact of climatic and atmospheric parameters on the diversification rate of plastid lineages. The estimated divergence times enabled us to reinterpret taxonomic classification of controversial fossils.

## Introduction

Sometime around 2.45 billion years ago (Bya), the Earth’s atmosphere started to change due to the accumulation of molecular oxygen. This transition, called the Great Oxidation Event, was a byproduct of sun-powered and water-splitting reaction of photosynthesis performed by cyanobacteria. Undoubtedly, cyanobacteria have fundamentally altered the conditions of life on our planet, but they did more than evolve oxygenic photosynthesis, they also passed this ability to eukaryotes in the process of cyanobacterial endosymbiosis (1).

The first cyanobacterial endosymbiosis, i.e. the engulfment and integration of a cyanobacterium into the eukaryotic cell, involved a *Gloeomargarita*-like species and an unknown protist (2). This endosymbiotic merger gave rise to three photosynthetic lineages of Archaeplastida: *Glaucophyta* (glaucophytes), *Rhodophyta* (red algae) and Chloroplastida (green algae and land plants). Their cyanobacteria-derived photosynthetic organelles are called respectively: muroplasts, rhodoplasts and chloroplasts, and collectively primary plastids (3). Two primary plastid-containing lineages, red and green algae, subsequently triggered an evolutionary radiation via a series of eukaryote-to-eukaryote endosymbioses that resulted in a multitude of secondary and higher-order plastids present in: cryptophytes, haptophytes, euglenids, chlorarachniophytes, dinoflagellates, stramenopiles and even parasitic apicomplexans (4).

The second cyanobacterial endosymbiosis involved the ancestor of all photosynthetic *Paulinella* species: *P. chromatophora, P. micropora* and *P. longichromatophora*. The *Paulinella* genus, classified within the evolutionary lineage of Rhizaria, comprises both photoautotrophic and heterotrophic species of testate filose amoeba (5, 6). The most important feature of the photosynthetic *Paulinella* are blue-green bodies called chromatophores that probably evolved from a picocyanobacterium of the *Prochlorococcus*/*Synechococcus*/*Cyanobium* clade. Compared to primary plastids, chromatophores represent cyanobacteria-derived organelles at an earlier stage of endosymbiosis, and they are the sole reason why *Paulinella* is so important from the evolutionary point of view (7, 8).

While the emergence of photosynthetic *Paulinella* is an evolutionary curiosity, the emergence and radiation of Archaeplastida have substantially shaped the Earth’s biosphere. Archaeplastidans have participated in carbon dioxide fixation, oxygen production and other global biogeochemical cycles for hundreds of millions of years, but they also donated their plastids to many protist groups thereby further spreading photosynthesis to multiple branches of the tree of life (4, 9). Due to the oxygen increase in water and atmosphere, Archaeplastida played a critical role in the evolution of new high-energy consuming multicellular organisms (9, 10). They are also the main primary producers in many ecosystems and the largest component of biomass on Earth (∼80%) (11). Around 500 million years ago (Mya), their representatives invaded land and made it suitable for animal colonization (12).

Given the global importance of Archaeplastida, it is not surprising that the group has been extensively studied for many decades. However, there are still some fundamental inconsistencies around their evolution, such as the diversification time and branching order among the Archaeplastida lineages, or even the number of cyanobacterial endosymbioses that triggered their evolution. These inconsistencies result from the fact that (i) archaeplastidans poorly preserve in a fossil state, (ii) the phylogenetic signal contained in their genomes has eroded to a large extent, and (iii) glaucophytes have been notoriously underrepresented in phylogenetic and molecular clock studies (13).

In order to bring us closer to resolving the contentious issues about Archaeplastida evolution, researchers have constructed their phylogenetic and molecular clock trees using various molecular markers, species, calibration points, phylogenetic and molecular clock methods (Tab. S1). As a result, they obtained a multitude of chronograms that are difficult to compare because they represent different tree topologies, i.e. *Glaucophyta* or *Rhodophyta* or Chloroplastida as the first branching Archaeplastida lineage. However, these chronograms do inform us about the diversification times of extant members of Archaeplastida though the dates vary considerably among the authors, and the differences sometimes even exceed one billion year (Tab. S1). Generally, most researchers agree that Archaeplastida evolved prior to ∼1.6 Bya, *Rhodophyta* prior to ∼1.2 Bya and Chloroplastida prior to ∼1.0 Bya; these ages are also supported by the oldest fossils of multicellular red alga *Rafatazmia chitrakootensis* (∼1.6 Bya) (14) and green alga *Proterocladus antiquus* (∼1.0 Bya) (15).

In this study, we investigate the diversification time and branching order among the photosynthetic organelles of Archaeplastida, some of their secondary descendants and for the first time perform multi-clock studies with all *Paulinella* photosynthetic species. We constructed 11 phylogenies and 18 chronograms using the largest number yet of plastid-encoded proteins. The obtained molecular clocks were also used to verify tentatively assigned cyanobacteria and Archaeplastida microfossils. In order to check the influence of calibration fossils on chronograms, we tested three calibration sets, including one with the recently discovered *R. chitrakootensis* (14) and *P. antiquus* (15). For each calibration set, we evaluated the discrepancies in age estimations by different clocks. Moreover, based on all chronograms, we also assessed the putative impact of volcanic activity, global and oceanic temperature as well as atmospheric concentration of carbon dioxide and oxygen on the diversification rate of photosynthetic organisms.

## Results and discussion

### Phylogenetic analyses

In MrBayes (16) and Beast (17), the inference of phylogenies and node ages was performed simultaneously while in PhyloBayes (18) the chronograms were calculated using fixed trees from IQ-TREE (19) (Fig. 1, S1-S18). Additionally, we used RAxML (20) to verify the tree topology with one additional maximum likelihood method (Fig. S19). Our analyses strongly support the monophyly of all plastids, which is common in phylogenies based on plastid markers (13), but with the exclusion of much recently and independently acquired chromatophores of *Paulinella* species. Both photosynthetic organelles of Archaeplastida and *Paulinella* diverged from early-branching freshwater cyanobacteria. The former groups with *Gloeomargarita lithophora* and the latter with *Cyanobium gracile* as their most closely related cyanobacteria relatives, consistent with previous studies (2, 8).

**Figure 1.**
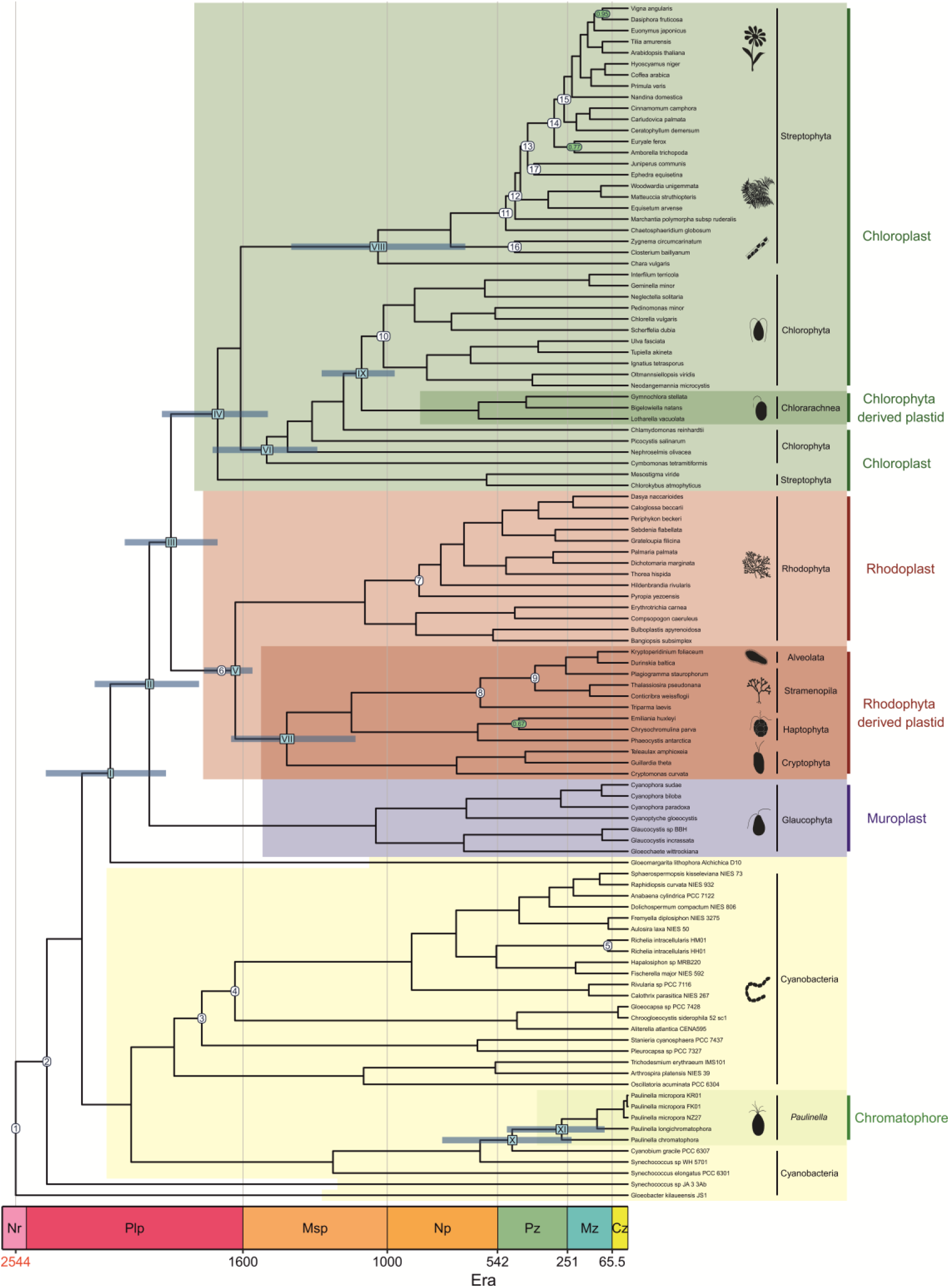
Time-calibrated phylogeny of photosynthetic organelles and cyanobacteria. The tree was inferred with Beast under UCLNR model and calibrated with data set C3 (Table 1). Arabic numerals in white circles indicate the calibration constraints. The roman numerals in blue rectangles mark key evolutionary events for plastids discussed in the article. At these nodes, there are blue bars representing 95% credibility intervals of the node age. The nodes supported with posterior probability lower than one are indicated with green circles.

All calculated 11 phylogenies favoured the glaucophyte-first hypothesis, i.e. *Glaucophyta* as the first branching lineage to the sister-group of *Rhodophyta* and Chloroplastida; nine Bayesian trees strongly supported this topology but the maximum likelihood methods did not (Fig. 1, S1-19). Our results agree with the greatest number of cyanobacterial characteristics preserved in muroplasts compared with rhodoplasts and chloroplasts, e.g. the presence of peptidoglycan and carboxysomes (13). However, morphological conservatism does not have to necessarily mean that *Glaucophyta* diverged first because the other Archaeplastida lineages could have simply lost certain ancestral traits independently. In accordance with this, previous phylogenetic analyses based on plastid markers support all the possible evolutionary models, which are in descending order of support: *Glaucophyta-*, Chloroplastida- or *Rhodophyta*-first hypothesis (1, 2, 13, 21, 22). Interestingly, the trees built on nuclear markers that do support the monophyly of Archaeplastida prefer either the *Glaucophyta* or *Rhodophyta* as the first branching lineage (13, 23–28). There is also generally greater support for the former scenario (13, 28) but, on the other hand, the most recent studies seem to favor the latter (23–27).

Similarly to Sánchez-Baracaldo et al. (29), *Chlorokybus* and *Mesostigma* broke up the monophyly of *Streptophyta* in all our phylogenies, and they were placed at the base of Chloroplastida with strong or moderate support (Fig. 1, S1-19). Consequently, these green algae possibly branched before the split of *Streptophyta* and *Chlorophyta*. However, this position was questioned by Lemieux et al. (30). Although they recovered similar topology in the maximum likelihood trees on the set of 45 plastid proteins, their trees based on the same set but nucleotide sequences supported *Chlorokybus* and *Mesostigma* as the earliest-branching *Streptophyta*. Moreover, their further analyses of gene content and/or gene order, and trees based on the amino acid sequences enriched in slow-evolving sites do preserve the monophyly of *Streptophyta* with both these green algae species (30).

The most interesting result from our phylogenies is the fact that *Paulinella* groups with *C. gracile* instead of *Prochlorococcus* or *Synechococcus* species (Fig. 1, S1-19). This allows us to formulate a new scenario for chromatophore acquisition. Under this scenario, the ancestors of photosynthetic *Paulinella* dwelled in marine habitats and fed on common picocyanobacteria, i.e. *Prochlorococcus* and *Synechococcus*, similarly to extant heterotrophic *Paulinella* (31). We assume that there was a preference for small prey like picocyanobacteria because *Paulinella* cells are surrounded by an oval theca, and consequently, the food has to fit through a narrow terminal aperture (∼1 µm in *P. ovalis*) (32). After transition to a new brackish/freshwater habitat, there would be no exclusively marine *Prochlorococcus*, but instead freshwater *Cyanobium* and possibly also some *Synechococcus* spp. (33). As a result, *Paulinella* would switch to *Cyanobium* as the more accessible prey. Next, it would learn to keep *Cyanobium* for prolonged time, e.g. as food for later use, and simultaneously profit from undergoing photosynthesis by absorbing carbon compounds, and possibly other nutrients, without killing the cyanobacterial prey. The transport of photosynthetic products and the other nutrients could provide the selective pressure for forging the endosymbiotic bond. In the final stage of endosymbiotic transformation, the host cell would abandon phagotrophy for the full photoautotrophic existence. Interestingly, it has been shown that eutrophic habitats preferred by photosynthetic *Paulinella* do drive unicellular eukaryotes into specialized trophic strategies, i.e. either photoautotrophy or heterotrophy (34).

### Molecular clock analyses

In total, we calculated 18 chronograms (Fig. 1, S1-17), six for each of the three calibration sets: C1, C2 and C3 (Tab. 1). C3 represents our most up to date calibration set since it includes microfossils of *R. chitrakootensis* (14) and *P. antiquus* (15). The other sets (C1 and C2) were used for comparative studies to investigate the impact of calibration constraints on molecular clocks (see below). The estimated ages are presented as ranges of mean dates from all six molecular clocks: UCLNR, IGR, TK02, CIR, LN and UGAM, unless stated otherwise (for details, see Materials and methods). We discuss only selected results concerning the evolution of major photosynthetic lineages though more datings are also included in the section: Verification of tentatively assigned microfossils in Supplementary Information.

**Table 1.**
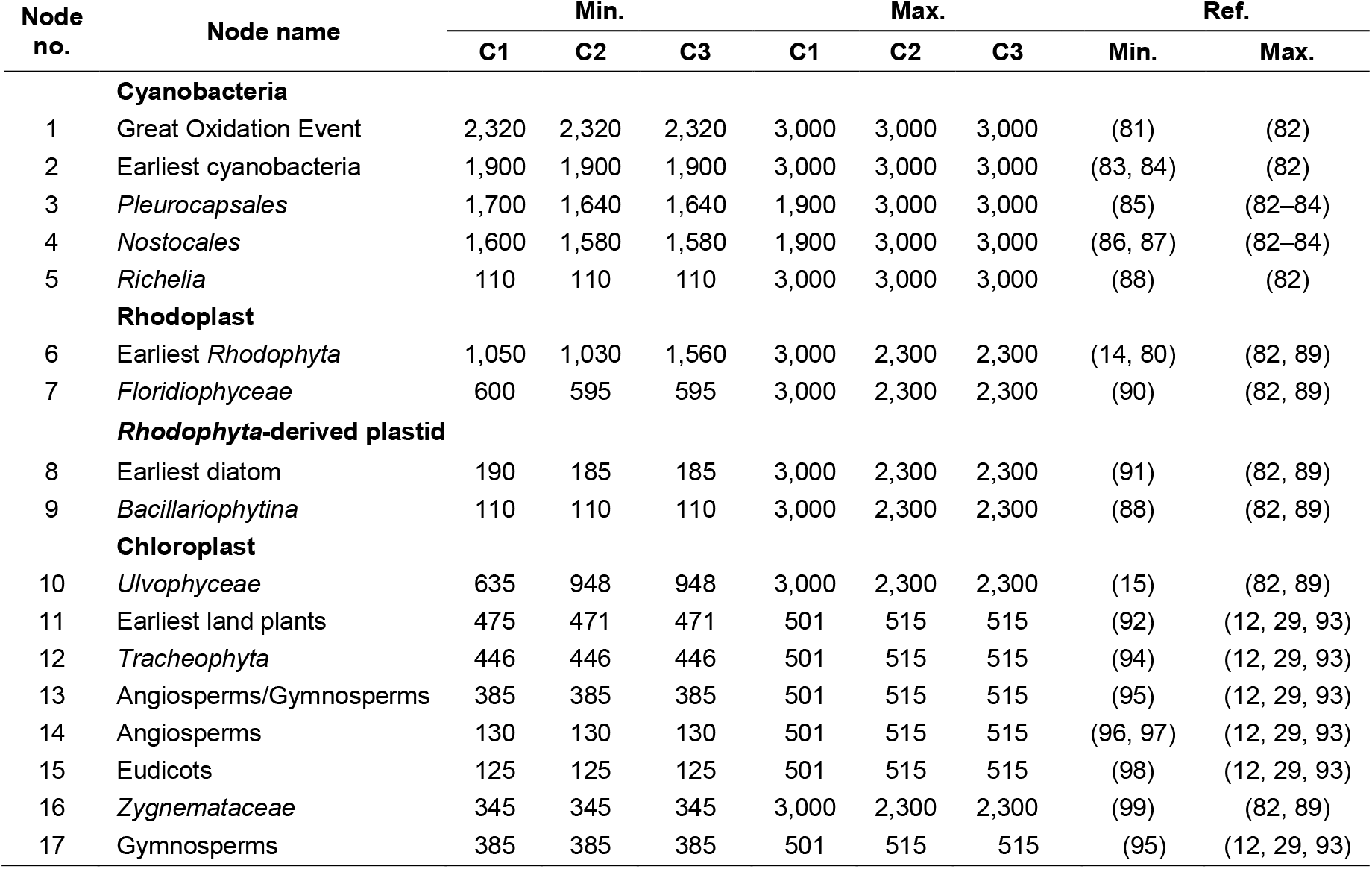
Calibration constraints for dating plastid evolution. A detailed description of each calibration point is provided in Tab. S2. C1, C2 and C3 represent the first, second and third calibration set, respectively. C1 was similar to the calibration strategy by Sánchez-Baracaldo et al. (29). C2 and C3 differ from C1 over using the microfossil of *P. antiquus* to calibrate the minimum age for the node 10 (15). In turn, C2 vary from C3 in calibration for node 6, the former uses widely recognized *Bangiomorpha pubescens* (80) while the latter much older and recently discovered *R. chitrakootensis* (14). Further small differences between C1 and the other calibration sets result from the fact that for C2 and C3, we always applied the minimum or maximum constraints on nodes by selecting the lower or upper interval for microfossil dating, respectively.

| Node no. | Node name | Min. |  |  | Max. |  |  | Ref. |  |
| --- | --- | --- | --- | --- | --- | --- | --- | --- | --- |
|  |  | C1 | C2 | C3 | C1 | C2 | C3 | Min. | Max. |
|  | **Cyanobacteria** |  |  |  |  |  |  |  |  |
| 1 | Great Oxidation Event | 2,320 | 2,320 | 2,320 | 3,000 | 3,000 | 3,000 | (81) | (82) |
| 2 | Earliest cyanobacteria | 1,900 | 1,900 | 1,900 | 3,000 | 3,000 | 3,000 | (83, 84) | (82) |
| 3 | *Pleurocapsales* | 1,700 | 1,640 | 1,640 | 1,900 | 3,000 | 3,000 | (85) | (82–84) |
| 4 | *Nostocales* | 1,600 | 1,580 | 1,580 | 1,900 | 3,000 | 3,000 | (86, 87) | (82–84) |
| 5 | *Richelia* | 110 | 110 | 110 | 3,000 | 3,000 | 3,000 | (88) | (82) |
|  | **Rhodoplast** |  |  |  |  |  |  |  |  |
| 6 | Earliest *Rhodophyta* | 1,050 | 1,030 | 1,560 | 3,000 | 2,300 | 2,300 | (14, 80) | (82, 89) |
| 7 | *Floridiophyceae* | 600 | 595 | 595 | 3,000 | 2,300 | 2,300 | (90) | (82, 89) |
|  | ***Rhodophyta*-derived plastid** |  |  |  |  |  |  |  |  |
| 8 | Earliest diatom | 190 | 185 | 185 | 3,000 | 2,300 | 2,300 | (91) | (82, 89) |
| 9 | *Bacillariophytina* | 110 | 110 | 110 | 3,000 | 2,300 | 2,300 | (88) | (82, 89) |
|  | **Chloroplast** |  |  |  |  |  |  |  |  |
| 10 | *Ulvophyceae* | 635 | 948 | 948 | 3,000 | 2,300 | 2,300 | (15) | (82, 89) |
| 11 | Earliest land plants | 475 | 471 | 471 | 501 | 515 | 515 | (92) | (12, 29, 93) |
| 12 | *Tracheophyta* | 446 | 446 | 446 | 501 | 515 | 515 | (94) | (12, 29, 93) |
| 13 | Angiosperms/Gymnosperms | 385 | 385 | 385 | 501 | 515 | 515 | (95) | (12, 29, 93) |
| 14 | Angiosperms | 130 | 130 | 130 | 501 | 515 | 515 | (96, 97) | (12, 29, 93) |
| 15 | Eudicots | 125 | 125 | 125 | 501 | 515 | 515 | (98) | (12, 29, 93) |
| 16 | *Zygnemataceae* | 345 | 345 | 345 | 3,000 | 2,300 | 2,300 | (99) | (82, 89) |
| 17 | Gymnosperms | 385 | 385 | 385 | 501 | 515 | 515 | (95) | (12, 29, 93) |

According to our chronograms based on C3 (Table 1), the cyanobacterial ancestors of muroplasts, rhodoplasts and chloroplasts diverged from *G. lithophora* in the Paleoproterozoic Era between 2.2 and 2.0 Bya (Fig. 1, 2, S1-S5; Tab. S3). Soon after, they got involved in the endosymbiotic interaction with an unknown unicellular eukaryote, probably in a freshwater environment and most likely beginning as food (29). This endosymbiotic merger must have taken place before the divergence of the first Archaeplastida lineage. We indicate that the lineage in question was *Glaucophyta* and that the process occurred before 2.1 to 1.8 Bya (Fig. 1, 2, S1-S5; Tab. S3). Our age for the first cyanobacterial endosymbiosis is in line with the chronograms calculated on nuclear markers by Strassert et al. (23), plastid markers by Blank (35) and Sánchez-Baracaldo et al. (29), but much older than the other estimations summarised in Tab. S1.

**Figure 2.**
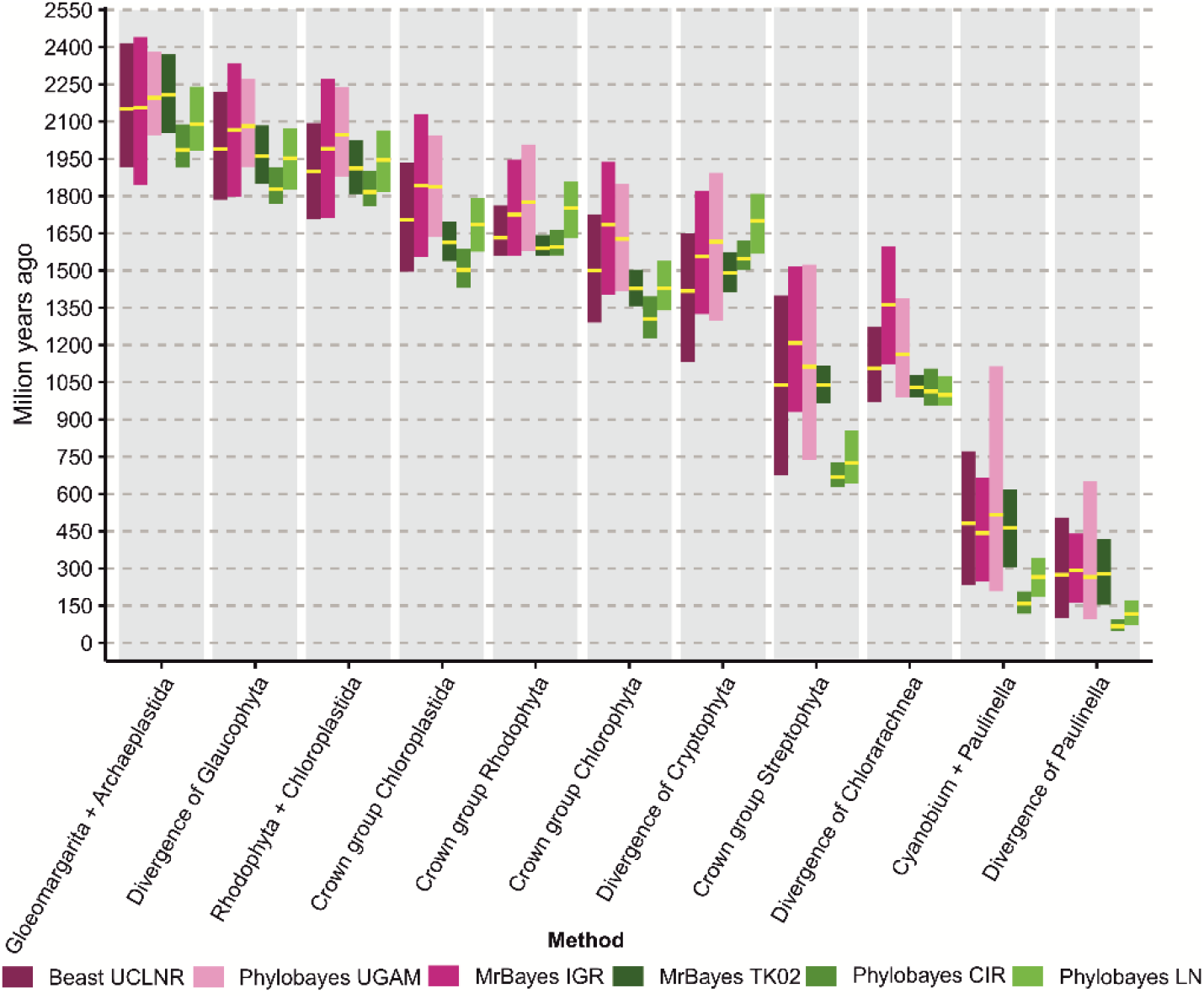
Comparison of molecular clock estimations for key evolutionary events for photosynthetic organisms discussed in the article. Purple and green colours mark uncorrelated and autocorrelated clocks, respectively. The yellow lines indicate the mean age and the bars 95% credibility intervals for the mean.

Following the emergence of glaucophytes, red and green algae diverged between 2.0 and 1.8 Bya (Fig. 1, 2, S1-S5; Tab. S3). Their first species were unicellular algae, but the multicellularity in Archaeplastida could have evolved as early as 1.9 Bya and surely before 1.6 Bya. The first date represents the age of controversial fossil of *Grypania* (for details, see Supplementary Information), possibly an early red or green multicellular alga (36), and the latter *R. chitrakootensis* (14). According to our clocks, the crown groups of Chloroplastida and *Rhodophyta* were both established in the late Paleoproterozoic Era, between 1.8 and 1.6 Bya; only the CIR model inferred ∼1.5 billion years (Byr) for Chloroplastida. This time range is much older than the previous estimations (Tab. S1) that indicated the Mesoproterozoic and for extant Chloroplastida also the Neoproterozoic Era as the period of origin (Fig. 1, 2, S1-S5; Tab. S3). The only exceptions were datings by Blank (35) and Strassert et al. (23) for Chloroplastida and *Rhodophyta*, respectively; they anchored the crown groups in the late Paleoproterozoic as well (Tab. S1).

The Chloroplastida lineage is made up of two large clades: *Chlorophyta* and *Streptophyta*. The former includes mainly marine green algae and the latter freshwater green algae (charophytes) and all land plants. We estimated the age for the crown group of *Chlorophyta* between 1.7 and 1.3 Bya (Fig. 1, 2, S1-S5; Tab. S3), and it was similar to Blank’s (35) results, but much older than the other molecular clock indications (Tab. S1). Interestingly, the extant streptophytes seem to be younger according to our molecular clocks compared with recent reports (Tab. S1), except for Sánchez-Baracaldo et al. (29). We inferred that they evolved in the late Mesoproterozoic, between 1.2 and 1.0 Bya (Fig. 1, 2, S1, S2, S5; Tab. S3). However, this time range does not take into account the mean dates calculated by CIR (Fig. S3) and LN (Fig. S4) clocks (∼700 Myr), which differed strongly from our other models.

Single-celled green algae belonging to *Chlorophyta*, class *Ulvophyceae*, got involved in an endosymbiotic interaction with a cercozoan amoeba resulting in a new photosynthetic lineage *Chlorarachnea* (37). Chlorarachniophytes, e.g. *Bigelowiella*, are best known for their nucleomorph, a vestigial nucleus preserved in the reduced ulvophycean cell that now represents a secondary plastid. According to our studies, this complex plastid split from their ancestral green algae between 1.1 and 1.0 Bya (Fig. 1, 2, S2-S5; Tab. S3), only the IGR model (Fig. S1) indicated an earlier mean age (1.36 Byr). These results are consistent with nuclear marker-based clock by Parfrey et al. (38), dating this event to about 1.0 Bya, but not with estimations based on plastid proteins by Jackson et al. (37), assuming the mean date ∼600 Mya.

Nucleomorph is not only characteristic of *Chlorarachnea* but also *Cryptophyta*; however, both organelles were acquired independently from green and red algae, respectively. Contrary to chlorarachniophytes, the red alga acquisition triggered plastid transfers to other protist lineages in tertiary or indirectly in higher order endosymbioses, e.g. to stramenopiles, haptophytes and dinoflagellates (23). The up to date scenarios for evolution of red alga-derived plastids mostly agree that there was a single secondary endosymbiosis beginning with cryptophytes and a few subsequent serial plastid acquisitions (see 23 and citations therein). Our results indicate that the process started between 1.7 and 1.4 Bya (Fig. 1, 2, S1-S5; Tab. S3), which is similar to Strassert et al. (23) estimation.

The minimum age for the second cyanobacterial endosymbiosis amounting to about 60 Mya was first proposed by Nowack et al. (7). They assumed that the rate of pseudogene disintegration requiring from 40 to 60 Myr in the endosymbiotic bacteria *Buchnera aphidicola* is comparable to that of *Paulinella* in chromatophores. Subsequent molecular clock studies based on 18S rRNA with the UCLNR model by Delaye et al. (39) and also Lhee et al. (40) estimated that photosynthetic *Paulinella* species diverged from their heterotrophic relatives 141-94 Mya and 193-64 Mya, respectively. In turn, Sánchez-Baracaldo et al. (29) based on plastid markers calculated the split of *P. chromatophora* from the *Synechococcus*/*Cyanobium* clade between 634 Mya and 350 Mya.

We are the first to present multi-clock studies that include all known *Paulinella* photosynthetic species. According to our four clocks, cyanobacterial ancestors of chromatophores diverged from *C. gracile* in the early Paleozoic Era between 516 and 443 Mya (Fig. 1, 2, S1, S2, S5; Tab. S3). These results are in line with Sánchez-Baracaldo et al. (29) estimations but we managed to narrow the upper and lower limits of the time range for about 100 Myr each. The second cyanobacterial endosymbiosis must have taken place before 292 to 266 Mya, i.e. before *P. chromatophora* separated from the other photosynthetic *Paulinella* species. It would mean that this endosymbiosis is much older than the other authors assumed. However, these time ranges do not include mean dates calculated by LN and CIR clocks, which strongly differed from our other models. LN and CIR chronograms indicate that *Paulinella* split from *C. gracile* between 266 and 158 Mya, respectively, and therefore acquired plastids between 118 and 67 Mya (Fig. 2, S3-S4; Tab. S3). In turn, these ages agree better with Delaye et al. (39) and Lhee et al. (40) datings.

### The impact of molecular clocks and calibration sets on age estimations

In order to evaluate the discrepancies in age estimations by different approaches, we compared (i) different calibration sets assuming the same molecular clock and (ii) a given calibration set with different molecular clocks.

The first comparison indicates that the data set C3 (Tab. 1) always provided older ages than the global mean for all the three calibration sets. The average percentage difference from the global mean per node *D* (see equation in Materials and methods) for C3 depended on the clock model and ranged from about 1% to 7% (Tab. 2). The ages estimated based on C1 were mostly younger than the global mean, from about 1% to 9%, and those based on C2 were in the middle in this respect (Tab. 1, 2). We also visualized these tendencies in Fig. S20 as pairwise comparisons of the calibration sets, and the differences between them were statistically significant (p < 2e-45). The largest average pairwise differences in ages per node *P* (see equation in Materials and methods) equal to 135 Myr and 121 Myr were observed for the comparison of C1 with C3 for clocks TK02 and LN, respectively, and they corresponded to the average percentage pairwise difference per node *P%* (see equation in Materials and methods) 17% and 14%. It is also worth mentioning that molecular dating by the UGAM model was least affected by the change in the calibration set (Tab. 2).

**Table 2.** The average percentage difference from the global mean per node *D* calculated for the three calibration sets assuming the same molecular clock model. The values should be compared in rows. U – states for uncorrelated model and A – for autocorrelated model.

| **Clock** | **Calibration set** |  |  |
| --- | --- | --- | --- |
|  | **C1** | **C2** | **C3** |
| Beast UCLNR (U) | -3.57 | -0.33 | 3.90 |
| MrBayes IGR (U) | -1.30 | -2.47 | 3.77 |
| MrBayes TK02 (A) | -9.30 | 1.90 | 7.40 |
| Phylobayes CIR (A) | -2.06 | 0.05 | 2.01 |
| Phylobayes LN (A) | -8.32 | 3.24 | 5.08 |
| Phylobayes UGAM (U) | -0.84 | 0.14 | 0.69 |

Much greater differences could be noticed for the second comparison, i.e. of different molecular clocks with the same calibration set (Tab. 3). For each calibration set, the LN model estimated on average from 13% to 20% older ages than the mean calculated for all the six chronograms. Quite old ages were also obtained for the IGR clock, from 6.5% to 13% greater than the global mean. The younger ages were produced by UCLNR, and they were about 18% smaller than the global mean. UGAM and CIR also provided younger ages but the results of TK02 depended on the calibration set. The pairwise comparison between the clocks is presented in Fig. S21, and the differences in the estimated ages are statistically significant for the comparison of LN and UCLNR with the other five models (p < 6.5e-07), for the comparisons of UGAM-CIR (p = 0.041) and UGAM-IGR (p = 0.019). As expected from the global comparison, we found the biggest average pairwise difference in ages per node *P* between the LN and UCLNR clocks. The ages were on average 208 Myr, 273 Myr and 282 Myr older for LN than UCLNR for C1, C2 and C3 sets, respectively, which corresponds to the mean percentage pairwise difference per node *P%* 25%, 33% and 31%.

**Table 3.** The average percentage difference from the global mean per node *D* calculated for the six molecular clocks assuming the same calibration set. The values should be compared in columns. U – states for uncorrelated model and A – for autocorrelated model.

| Clock | Calibration set |  |  |
| --- | --- | --- | --- |
|  | C1 | C2 | C3 |
| Beast UCLNR (U) | -17.93 | -18.58 | -18.05 |
| MrBayes IGR (U) | 13.16 | 6.51 | 9.79 |
| MrBayes TK02 (A) | -5.61 | 0.49 | 2.21 |
| Phylobayes CIR (A) | -0.15 | -3.19 | -4.29 |
| Phylobayes LN (A) | 13.03 | 20.96 | 19.14 |
| Phylobayes UGAM (U) | -2.50 | -6.19 | -8.80 |

We also performed multidimensional scaling to visualize the similarities and differences indicated above for all the combinations of calibration sets and molecular clocks (Fig. 3). This analysis illustrates that there are clearly greater differences in the age estimations between molecular clocks compared to calibration sets. In most cases, the calibration sets were grouped together independently of the clock model applied, but the clocks were generally scattered across the plot.

**Figure 3.**
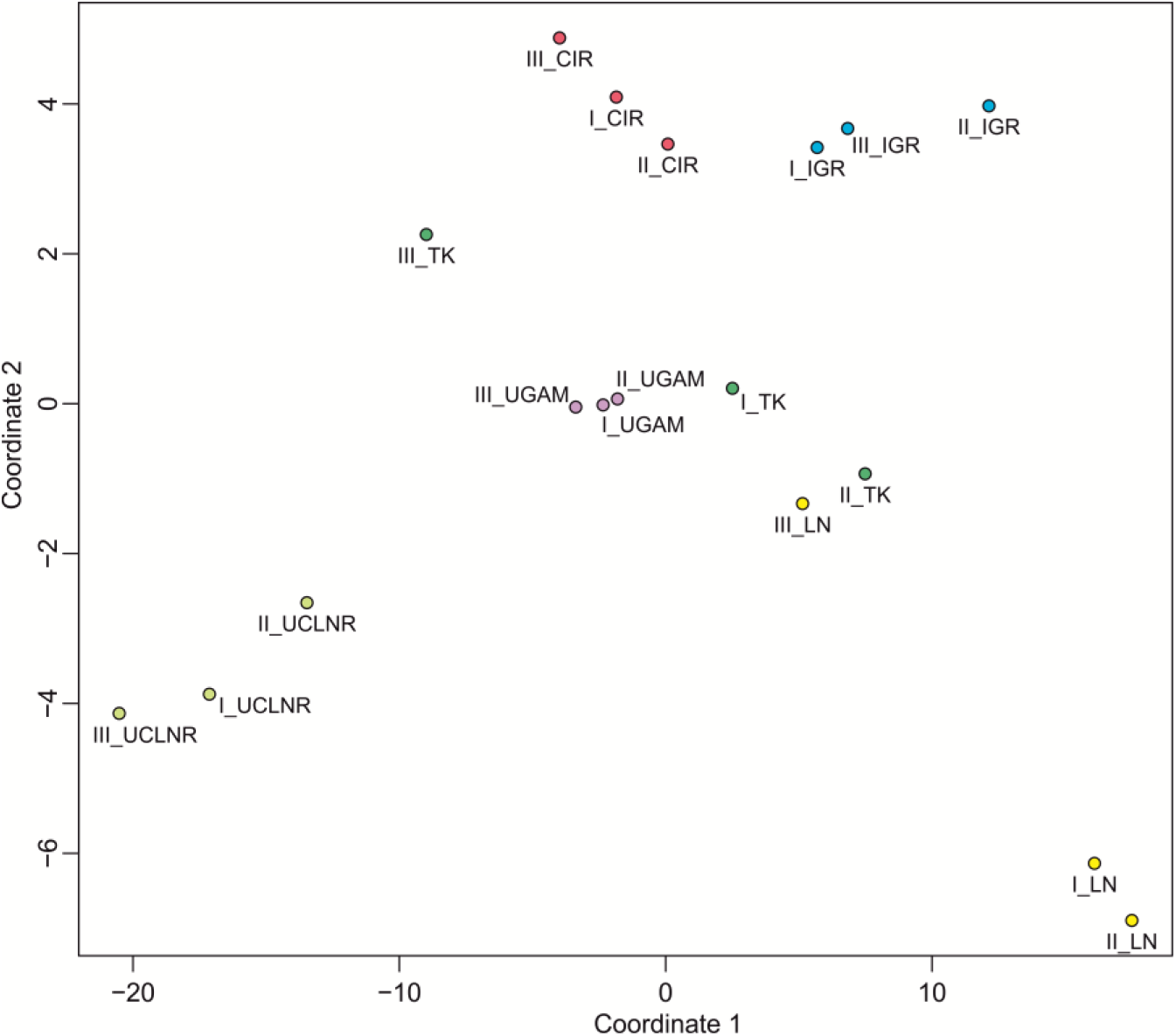
Visualization of similarities and differences in age estimation for all combinations of calibration sets and molecular clocks in multidimensional scaling. The three calibration sets with the UCLNR clock (light green dots) and the set C2 and C3 with the LN clock (yellow dots) represent the most distant points because they calculated the most extreme age estimations in our analyses, the youngest and the oldest respectively. The other combinations of clocks and sets, depicted as dots of appropriate colour, are located between them since they provided more moderate datings.

### Impact of climatic and atmospheric parameters on diversification rate

The obtained chronograms (Fig. 1, S1-17) were used to estimate the diversification rates of photosynthetic organisms and their correlation with various climatic and atmospheric parameters. We took into account the global and oceanic temperature as well as the atmospheric concentration of carbon dioxide and oxygen (Tab. S9). The volcanic activity was approximated by the area of Large Igneous Provinces (Tab. S9). We also calculated the mean diversification rate during six main glaciations and in the warmer periods between them (Tab. S10).

All statistically significant Spearman correlation coefficients (ρ) calculated for various combinations of the diversification rates with temperature and CO_2_ were positive (Fig. 4). The mean ρ was 0.54 and 0.61, and for some cases this coefficient reached 0.7 and 0.93, respectively. In accordance with that, we found a significantly higher diversification rate in warmer periods, compared to glaciations (p-value = 0.00006). The mean ratio between the warm periods and glaciations for the diversification rate was 1.41, and it amounted to 2.35 in some extreme cases.

**Figure 4.**
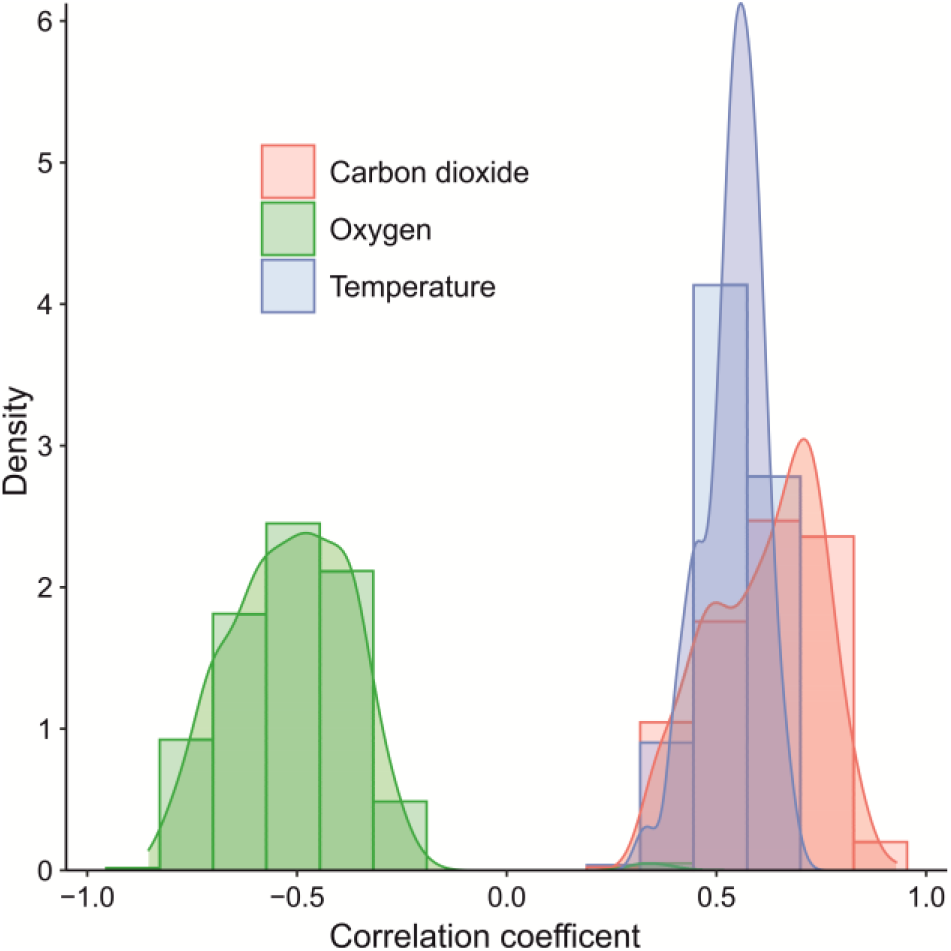
Distribution of statistically significant Spearman correlation coefficients calculated for various combinations of the diversification rates with many parameters of temperature as well carbon dioxide and oxygen concentration.

Our results indicate that the diversification rate of photosynthetic organisms significantly rose in warmer times under higher CO_2_ concentrations. Significantly, these conditions have already been reported to have a positive impact on photosynthesis and the global growth and reproduction of various alga and plant species (41–44). Additionally, the increase in CO_2_ may reduce the stomatal conductance and transpiration in plants and, therefore, alleviate the negative influence of drought and improve the use of nutrients and water (45). A similar beneficial effect might produce the rising temperature, but only to some extent. It results in higher enzyme activity (46) that, in turn, is also associated with the stimulation of cell growth and proliferation (47–51).

The positive impact of the carbon dioxide and temperature can be further enhanced by their synergy (49, 52). For example, the influence of elevated CO_2_ on photosynthesis is stronger at higher temperatures due to the suppression of photorespiration, a competitive reaction to photosynthesis (53). Moreover, higher concentration of CO_2_ can shift the optimum temperature for plant photosynthesis, growth and development to greater values (53–55).

As a consequence of the positive effect of CO_2_ and temperature, the population size of photosynthetic organisms would increase, and trigger higher speciation rates in accordance with the unified neutral theory of biodiversity (56). The theory postulates that ‘the number of new species arising per unit time is a function of the total number of individuals in a metacommunity’. Warmer temperatures are also responsible for boosting metabolism, higher mutation rates and shorter generation times, which may accelerate speciation as well (57, 58). Elevated temperatures were shown to increase the selection strength on genome-wide polymorphism causing more rapid evolution and diversification in warmer regions (59). Accordingly, there is increasing species richness of pteridophytes and seed plants with growing temperature (60, 61).

Our findings are in agreement with Barrett and Willis (62); they postulated that the major plant groups emerged at times of increasing CO_2_. More detailed analyses showed that CO_2_ concentration was strongly and positively correlated with gymnosperm and pteridophyte speciation (63). In another study, the diversity in neotropical wet forests was also positively related with CO_2_ over long timescales during the Cenozoic (64). Moreover, Fiz-Palacios et al. (65) found that the majority of shifts in the diversification rate of angiosperms, ferns, and mosses coincided with warm climates.

In contrast to the positive effect of CO_2_ and temperature, the significant correlations between the diversification rate and O_2_ concentration were in 99.4% cases negative (Fig. 4). The mean ρ was -0.51 and in an extreme case it amounted to -0.85. This can be associated with an adverse role of oxygen in photosynthesis. O_2_ is a competitive inhibitor of CO_2_ fixation by ribulose-1,5-bisphosphate carboxylase/oxygenase (66) and stimulates photorespiration (67). A negative influence of oxygen could also be related to fueling fire on land biota and it was included in some paleoclimatic models (68).

A weak negative correlation was also noticed between the diversification rate and the volcanic activity (data not shown). Regardless of the measure used (for details, see Materials and methods), the mean ρ was -0.29 for statistically significant cases. Considering a long-term influence, the negative effect of volcanos could be related to blocking sunlight and climate cooling by ash and dust as well as to droplets of sulfuric acid, which decrease photosynthesis activity (69, 70). Covering leaves and their stomata by the ash and dust can also disturb respiration processes, whereas acidic rains can cause their direct damage. However, volcanos are important sources of CO_2_, a substrate to photosynthesis. Thus, this two-sided influence can cause that the correlation coefficient is not very high but the negative effect apparently prevails.

### Concluding Remarks

According to our phylogenetic and molecular clock analyses, the primary plastids evolved prior to 2.1 to 1.8 Bya, i.e. before the first Archaeplastida lineage, *Glaucophyta*, diverged. Similarly to the primary plastids, *Paulinella* chromatophores evolved in low salinity habitats because they group in all our trees with the freshwater picocyanobacterium *C. gracile*. This relatively recent endosymbiosis took place possibly before 292 and 266 Mya though two of six clocks indicated younger ages. Following our chronograms, the red algae secondary endosymbioses started with cryptophytes between 1.7 and 1.4 Bya, whereas chlorarachniophytes acquired their green alga-derived plastids between 1.1 and 1.0 Bya (Fig. 1, 2). Generally, the age estimations for our key evolutionary events concerning photosynthetic organelles were older compared to previous reports (Tab. 1S), and at least partially due to the inclusion into our calibration set of new microfossils: *R. chitrakootensis* (14) and *P. antiquus* (15); they represent the oldest-known red and green alga specimens, respectively. The diversification rate of the investigated photosynthetic organisms was positively correlated with temperature and carbon dioxide concentrations but negatively with oxygen levels and volcanic activity (Fig. 4). Our analyses of the impact of molecular clocks and calibration sets on the age estimations also indicate that there are clearly greater differences in the ages between the clocks compared to the calibration sets (Fig. 3). However, we do not see the dependence of uncorrelated and autocorrelated models with the estimation of younger and older ages, respectively, though some models seem to follow this rule.

## Materials and Methods

### Dataset preparation

We used carefully selected 30 conserved plastid-encoded proteins (Table S4) from five organisms: *Gloeomargarita lithophora, Cyanobium sp*., *Cyanophora paradoxa, Galdieria sulphuraria* and *Arabidopsis thaliana* for homologous search by PSI-BLAST (e-value: <0.001, number of iterations: 5, word size: 2) against amino acid sequences encoded by plastids, cyanobacteria and chromatophore genomes (71). Plastid proteins were retrieved from the NCBI reference sequence database (72) while cyanobacteria and chromatophore proteins from GenBank (73). The final set included 108 representatives of cyanobacteria and plastid-carrying eukaryotes (mainly archaeplastidans) and each contained the full number of plastid markers, with the exception of chlorarachniophytes that lacked CCSA protein. Each protein group was aligned independently in MAFFT v7.429 using a slow and accurate L-INS-i algorithm (74). The multiple sequence alignments were inspected in AliView (75) and the sites best suited for phylogenetic analyses were selected with TrimAl gappyout method (76). All 30 trimmed alignments were concatenated into the supermatrix of 9,823 aligned amino acid positions with SequenceMatrix 1.8 (77).

### Phylogenetic and molecular clock analyses

The phylogenetic trees were obtained based on: (i) the maximum likelihood method in IQ-TREE 2.0 (19) and RAxML 8.2.12 (20), and (ii) the Bayesian approach in MrBayes 3.2.7a (16) and Beast 2.6.0 (17) along with the molecular clock analyses. We considered all 30 potential partitions, represented by each protein group, in finding the best substitution models for the whole protein set. To estimate the best-fitted substitution models, we used ModelFinder (78) in IQ-TREE (19), and PartitionFinder (79) for RAxML (20), MrBayes (16) and Beast (17). They proposed 9 (Table S5) and 19 (Table S6-S8) partitions with individual substitution models, respectively. Branch support was inferred with 100 replicates of non-parametric bootstrap test for IQ-TREE (19) and RAxML (20), and estimation of posterior probability for MrBayes (16) and Beast (17).

We performed Bayesian molecular dating with two different relaxed molecular clock methods: autocorrelated and uncorrelated, which assume dependent or independent rate of evolution on adjacent branches of the phylogenetic tree, respectively. We implemented: autocorrelated log normal model (LN), Cox–Ingersoll–Ross model (CIR) and uncorrelated gamma multiplier model (UGAM) in PhyloBayes 4.1c (18); autocorrelated Thorne–Kishino 2002 model (TK02) and uncorrelated independent gamma rate model (IGR) in MrBayes (16) as well as uncorrelated lognormal model (UCLNR) in Beast (17). In total, we calculated 18 chronograms for three calibration sets and six molecular clock approaches.

In PhyloBayes, we used the fixed tree topology inferred in IQ-TREE and calculated each molecular clock under CAT-GTR+Γ model. We always selected *Gloeobacter kilaueensis* JS1 as an outgroup. For each molecular clock and calibration set, two replicate chains were run. Analyses were performed using hard bounded gamma-distributed prior for the root and uniform hard bounded priors for the rest calibration constraints. We selected burnin to get the maximum discrepancy lower than 0.3 and the minimum effective size greater than 50 in tracecomp. Mean dates were assessed by running readdiv with burnin selected for each analysis in tracecomp.

In Beast, we implemented Yule tree prior and exponential priors on calibration constraints. Tree samples were saved every 1000 iterations. The first 25% samples were discarded as burnin to get the effective sample size greater than 100 for all parameters calculated in Tracer 1.7. The final chronograms were obtained in Treeannotator using 25% burnin and mean heights.

In MrBayes, offset exponential priors were used for divergence times together with birth-death tree prior. Samples were saved every 100 iterations. To summarize the results, we used sump command discarding 25% of the first samples. We made sure that the potential scale reduction factor was close to 1 and the average effective sample size greater than 100 for all parameters. The final chronograms were obtained in Treeannotator using 25% burnin and mean heights.

### Comparison of molecular clocks and calibration sets

In order to assess a potential bias in the estimation of divergence times among molecular clocks and calibration sets, we calculated for each set the percentage difference from the mean age of all sets obtained for each node, and next the average differences across all nodes *D* according to the equation:

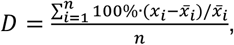

where: *x*_*i*_ is an age for the node *i*, 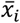 is a mean age of sets for the node *i* and *n* is the number of nodes. In the analysis we included 104 nodes and excluded three that were not shared in all tree topologies. The calculations were performed separately for: (i) different calibration sets assuming the same molecular clock as well as (ii) for each calibration set with different clocks.

Moreover, we calculated the mean pairwise difference in ages across all nodes between individual sets *P* according to the equation:

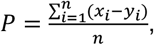

where: *x*_*i*_ is an age in the set *x* for the node *i, y*_*i*_ is an age in the set *y* for the node *i* and *n* is the number of nodes. Similarly, we calculated the mean percentage pairwise difference:

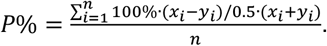

These values were used for multidimensional scaling (MDS) in R to compare age estimations under various assumptions in a graphical way. Statistical significance of differences between the compared sets was assessed using the paired Wilcoxon test. The resulted p-values were corrected using the Benjamini-Hochberg method.

### Comparison of diversification rate with climatic and atmospheric parameters

Based on the 18 chronograms, we calculated diversification rates within various periods using yuleWindow function in R. For each chronogram, we took into account the periods of 10, 20, 30, 40, 50, 60, 70, 80, 90 and 100 Myr as overlapping windows with the shift of the half of window length. The middle value of the window was ascribed to the calculated diversification rate.

The Spearman correlation coefficient and its significance were calculated between the values of diversification rate and various climatic/atmospheric parameters (Table S9). The parameter values for the same time points as for rates were obtained based on a fitted smoothing spline calculated by ss function in R. The p-values of the correlation coefficient were corrected using the Benjamini-Hochberg method to control the false discovery rate. Differences were considered significant when p-values were smaller than 0.05.

The relationship between the area of Large Igneous Provinces and geological age was expressed in two ways. In the first approach, the total area of Large Igneous Provinces was calculated in the window of 10 Myr from the available data. When the age range was given, the mean age was assumed for the data. In the second approach, the area per 1 Myr was calculated, when the age range was given, and next the sum was calculated in the period of 10 Myr. Finally, the middle value of the window was included in the relationship.

The mean diversification rate for 18 chronograms was calculated in the periods of six main glaciations and interglaciations (Table S10). The final results were presented as the ratio between the latter and the former rate values. The significance of the difference between these values was assessed in the unpaired Wilcoxon-Mann-Whitney test. All statistical analyses were performed in R.

## Acknowledgments

We are grateful to Richard Ernst for providing data about large igneous province records, and to Giada N Arney, David Catling, Joshua Krissansen-Totton and Dana Royer for paleoclimatic data and their useful explanations. This work was supported by the National Science Centre grant 2017/26/D/NZ8/00444 to P.G., the National Science Centre grant 2018/31/N/NZ2/01338 to K.S. and the National Science Centre grant 2019/35/N/NZ8/03366 to F.P.

## Notes

**Competing Interest Statement:** The authors declare no conflict of interest.

### Competing Interest Statement

The authors have declared no competing interest.

