## Supplementary Information for "Dating the photosynthetic organelle evolution in Archaeplastida, *Paulinella* and secondary-plastid bearing lineages"

**This PDF file includes:**

Supplementary text
Figures S1 to S21
Tables S1 to S10
SI References

### Supplementary Information Text

**Verification of tentatively assigned microfossils.** In the light of the obtained chronograms based on the calibration set C3, we were able to verify taxonomic affiliation of tentatively and controversially assigned fossils. Our estimations indicate that the main extant cyanobacterial lineages diverged between 2.5 and 1.6 Bya (Fig. 1, S1, S3-S5); only TK02 model indicated an older mean dating (2.74 Bya) (Fig. S2). Consequently, the assignment of individual Archaean microfossils to contemporary classes or orders of cyanobacteria is not substantiated based on our results. Actually, many of these specimens are the subject of hot debate because they may represent other bacteria groups or are considered not to be organismal remains at all (1). Our chronograms support the assignment of *Oscillatoriopsis* (2.2 - 1.8 Bya, Australia (2)) but not *Siphonophycus* (2.5 - 2.3 Bya, South Africa (3)) to *Oscillatoriales* because *Oscillatoriales* diverged between 2.1 and 1.8 Bya (Fig. 1, S1-S5). Moreover, *Eoentophysalis belcherensis*, the oldest microfossil interpreted with certainty as a cyanobacterium (~1.9 Bya, Canada), does not represent the contemporary lineage of *Chroococcales*; according to our results this order together with other coccoidal cyanobacteria (*Chroococcidiopsidales*) evolved between 1.8 and 1.6 Bya (4) (Fig. 1, S1-S5).

An interesting but controversial fossil represents *Grypania*, with the oldest occurrence described from 1.9 Byr-old formations in Michigan, USA (5, 6). *Grypania* has been commonly interpreted as a macroalga but alternatively named: a pseudofossil, trace fossil, cyanobacterium or even a metazoan (7). Its age falls well within the estimated time range for the split of the Archaeplastida lineages, i.e. between 2.1 and 1.8 Bya (Fig. 1, 2, S1-S5). Accordingly, *Grypania* might represent an early green or red alga though it would mean that the multicellularity evolved rather early in the archaeplastidian evolution.

*Bangiomorpha* is another example of an interesting microfossil and until recently it has been considered to be the oldest red alga specimen (~1.0 Bya, Canada (8)). Our chronograms indicate that it could be directly related to extant representatives of the order *Bangiales*, class *Bangiophyceae*, as morphological data suggest (8). We estimated the age for this group between 1.4 and 1.0 Bya (Fig. S1-S5); only the Beast model indicated a younger mean dating (868 Mya, Fig. 1).

According to our analyses, some acritarchs, i.e. unicellular organic-walled microfossils of Proterozoic era, described as *Dictyosphaera-Shuiyousphaeridium*, *Gigantosphaeridium* and *Leiosphaeridia*, could belong to the Chloroplastida as interpreted by some authors e.g. Agić et al. (9). Their oldest representatives were found in 1.7 - 1.4 Byr-old deposits in China (9), which is more or less at the time of the diversification of extant green algae in our chronograms (1.84 - 1.55 Bya) (Fig. 1, S2-S5). Moreover, our results also support the classification of some acritarchs to prasinophytes (9, 10). Some of the oldest fossils of such type are *Simia* (1.7 - 1.4 Mya, China), *Valeria* (1.7 - 1.4 Bya, China), *Tasmanites* (~1.6 Bya, India) and *Pterospermella* (1.3 - 1.2 Bya, Greenland); their age corresponds well with the divergence of prasinophytes in our study between 1.7 and 1.2 Bya (Fig. 1, S2-S5).

We estimated the split of the clade consisting of *Trebouxiophyceae*, *Ulvophyceae* and *Chlorophyceae* from prasinophytes between 1.5 and 1.2 Bya (Fig. 1, S2-S5). Consequently, the assignment to this group of *Osculosphaera*, *Trachyhystrichosphaera* and *Vandalosphaeridium* dated to 1.25 Bya could be correct (11). However, *Tappania plana* (1.7 - 1.4 Bya, China) probably represents a fungus, especially taking into account that only two models: IGR (Fig. S1) and UGAM (Fig. S5) indicated datings older than 1.4 Bya (9). *Leiosphaeridia gorda*, *Cerebrosphaera*, *Culcitulisphaera* and *Lanulatisphaera* presumably belong to *Chlorophyceae* (11, 12), and their oldest remains dated ~1.0 Bya from Canada and 800 - 750 Mya from Kazakhstan, Sweden and Spitsbergen do agree with the divergence of *Chlorophyceae* in our analyses estimated between 1.4 and 1.0 Bya (Fig. 1, S2-S5). The chronograms we obtained also support that *Archaeoclada*, *Variaclada* and *Eoprotoderma* described from ~1.0 Byr-old formations in Siberia (11, 13) belong to the *Ulvophyceae* class that evolved between 1.27 and 1.0 Bya (Fig. 1, S2-S5). However, the classification of *Spiromorpha* (1.7 - 1.4 Bya, China) to *Zygnematophyceae* seems inconsistent with our estimations because our results indicate that *Zygnematophyceae* separated only between 941 and 551 Mya (Fig. 1, S2-S5).

We also calculated the divergence of land plants (embryophytes) and the bryophyte-tracheophyte clade to 508 - 514 Mya and 479 - 462 Mya, respectively; only IGR (831 Mya, 618 Mya) and TK02 (660 Mya, 569 Mya) models indicated older mean datings (Fig. 1, S2-S5). Our estimations allow us to assume that the ~510 Myr-old spores from Arizona and Tennessee in USA (14) and ~480-Myr-old spores from Australia (15) could indeed represent the early land plants, and the ~460 Myr-old trilete spores from Sweden the vascular plants (16).

**Table S1.** Comparison of molecular clock estimates from various studies for the divergence of the crown groups of Archaeplastida.

| | **Strassert et al. 2021** (17) | | **Nie et al. 2020** (18) | | **Sánchez-Baracaldo et al. 2017** (19) | | **Yang et al. 2016** (20) | **Blank 2013** (21) | | **Parfrey et al. 2011** (22) | **Berney at al. 2006** (23) | **Yoon at al. 2004** (24) | | **Douzery et al. 2004** (25) | **Hedges et al. 2004** (26) |
|---|---|---|---|---|---|---|---|---|---|---|---|---|---|---|---|
| **Crown Group** | **MIN** | **MAX** | **MIN** | **MAX** | **MIN** | **MAX** | **MEAN** | **MIN** | **MAX** | **MEAN** | **MEAN** | **MIN** | **MAX** | **MEAN** | **MEAN** |
| Archaeplastida | 1807,29 | 2137,51 | 1644[bc] | 1644[bc] | 1900 | 1939 | 1693[c] | 2006 | 2274 | 1556[b] | 930 | 1535 | 1719 | 1017[bc] | 1428[c] |
| Glaucophyta | 681,61 | 1440,58 | - | - | 1695[a] | 1781[a] | - | 1729[a] | 1952[a] | 598[b] | - | 1535[a] | 1719[a] | - | - |
| Rhodophyta | 1606,29 | 1657,63 | 1264[b] | 1264[b] | 1060 | 1161 | 1511[b] | 1412 | 1504 | 1236[b] | 740[b] | 1349 | 1452 | 928[a] | 1428[ac] |
| Chloroplastida | 1064,73 | 1523,87 | 1162,9 | 1738,2 | 1254[b] | 1254[b] | - | 1597 | 1710 | 941[b] | 695[b] | - | - | 729 | 968 |
| Chlorophyta | 940,89 | 1376,12 | 1065,8 | 1428,3 | 1037 | 1092 | - | 1394 | 1599 | 851[b] | 616[b] | - | - | 729[a] | 968[a] |
| Streptophyta | 942,75 | 1394,45 | 1115,1 | 1575 | 880 | 999 | 482[b] | 1438 | 1607 | 739[b] | 604[b] | 646 | 792 | 450[b] | 707 |
| **Markers** | 320 nuclear-encoded proteins | | 81 chloroplast-encoded protein genes | | 26 plastid-encoded proteins | | nuclear-encoded EF2 protein, LSU and SSU rRNAs, mitochondrial-encoded Cox1 protein, and plastid-encoded RbcL, PsaA and PsbA proteins | plastid-encoded LSU/SSU rRNA genes | | 15 nuclear-encoded protein genes | plastid-encoded SSU rRNA gene | plastid-encoded SSU rRNA, *psaA*, *psaB*, *psbA*, *rbcL*, and *tufA* genes and PsaA, PsaB, PsbA, RbcL, and TufA proteins | | 129 nuclear-encoded proteinS | 50-74 nuclear-encoded proteins |

a – one species represents the crown group
b – age read from the tree
c – tree without any *Glaucophyta* representatives

**Table S2.** Description of calibration constraints used for dating plastid evolution.

| Age (Mya) | Type | Description | Ref(s). |
|---|---|---|---|
| 3,000 | Maximum | The oldest evidence for the presence of atmospheric oxygen capable of weathering rocks, based on oxidative chromium isotopes and other redox sensitive metals found in 3,000 Myr-old South African formations from the Pongola Supergroup (Nsuze palaeosol and Ijzermyn iron formation). This dating suggests that significant levels of atmospheric oxygen were present 600 million years before the Great Oxidation Event. | (27) |
| 2,320 | Minimum | The analysis of sulfur isotopes in the marine sediments from the Transvaal Supergroup (Rooihoogte and Timeball Hill formations) indicates that the oxygen level remained very low until about 2,320 Mya. This date represents the first major stable increase in atmospheric oxygen (the Great Oxidation Event) and is linked to the cyanobacterial oxygenic photosynthesis. | (28) |
| 2,300 | Maximum | Molecular clock analyses indicate this date as the origin of mitochondria (2300 - 1800 Mya). It exactly coincides with the Great Oxidation Event; after the rise of oxygen level, mitochondria and organisms with more than two or three cell types are presumed to have evolved. | (26) |
| 1,900 | Minimum | The microfossil of *Eoentophysalis belcherensis* is the oldest microfossil interpreted with certainty as a cyanobacterium. It was described from 1,900 Myr-old stromatolitic dolostones of the Belcher Supergroup (Kasegalik and McLeary formations) in Hudson Bay, Canada. *E. belcherensis* closely resembles extant cyanobacteria from the genus *Entophysalis*, order *Chroococcales*. | (29, 30) |
| 1,640/ 1,700 | Minimum | The microfossil of *Eohyella* represents another indisputable microfossil of cyanobacteria. It was described from 1,640/1,700 Myr-old stromatolites of the Changcheng Group (Dahongyu formation) in the Pangjiapu iron mine, China. *Eohyella* closely resembles extant cyanobacteria from the genus *Eohyella*, order *Pleurocapsales*. | (31) |
| 1,580/ 1,600 | Minimum | The microfossil of *Archaeoellipsoides* was described in samples from McArthur Basin, Australia. The date 1,589±3 represents the minimum age for McArthur group (Upper Balbirini Dolomite formation). The date 1,600 Mya refers to calibration point from Sánchez-Baracaldo et al. (19). | (32, 33) |
| 1,560 | Minimum | The microfossil of *Rafatazmia chitrakootensis* represents the first multicellular eukaryote of red alga origin. It was described from 1,650±89 Myr-old phosphatized stromatolitic microbialites of the Tirohan Dolomite, part of the Chitrakoot Formation (Semri Group), India. | (34) |
| 1,030/ 1,050 | Minimum | The microfossil of *Bangiomorpha pubescens* is indisputably recognized as a multicellular red alga. It was described from 1,047 +0.013/-0.017 Myr-old chert rocks of the Bylot Supergroup (Angmaat Formation) in Baffin Island, Canada. The date 1,050 Mya refers to the calibration point from Sánchez-Baracaldo et al. (19). *B. pubescens* resembles red algae in the order *Bangiales,* class *Bangiophyceae*. | (8) |
| 948 | Minimum | The microfossil of *Proterocladus antiquus* represents the first multicellular green alga. It was described from 1,056 - 947.8 Myr-old mudstones of the Xihe Group (Nanfen Formation), China. *P. antiquus* resembles extant algae in the order *Siphonocladales*, class *Ulvophyceae*. | (35) |
| 635 | Minimum | The date was used in molecular clock analyses by Verbruggen's et al. (36) as a maximum age constraint for siphonous algae (class *Ulvophyceae*), and by Sánchez-Baracaldo et al. (19) as a minimum age constraint for *Ulvophyceae*. | (36) |
| 595/600 | Minimum | Fossils of multicellular red algae that resemble species in the order *Corallinales*, class *Florideophyceae*. They were described from 599±4 Myr-old material of the Doushantuo Formation, China. The date 600 Mya refers to the calibration point from Sánchez-Baracaldo et al. (19). | (37) |
| 515 | Maximum | Molecular clock analyses indicate the date from 514.8 to 473.5 Mya as the origin of land plants. Moreover, at that time cryptospores were described from Cambrian Stage 4 mudstones of the Conasauga Group (Rome Formation), USA. | (38, 39) |
| 501 | Maximum | The date 501 Mya refers to the calibration point from Sánchez-Baracaldo et al. (19). | (19) |
| 471/475 | Minimum | Earliest known cryptospores described from 473 - 471 Myr-old samples of the Zanjon Formation, Argentina. The date 475 Mya refers to the calibration point from Sánchez-Baracaldo et al. (19). | (40) |
| 446 | Minimum | The microfossils of spores of vascular plants described from 446 Myr-old (Late Katian) material of the Qusaiba-1 corehole, Saudi Arabia. | (41) |
| 385 | Minimum | The microfossils of megaspores described from upper Eifelian and lower to middle Givetian samples of the Miastko 1 borehole, Poland. The megaspores resemble archaeopteridalean megaspores and megaspores from Carboniferous gymnospers. | (42) |
| 345 | Minimum | The microfossils of *Zygnemataceae* (zygospores) described from Tournaisian samples collected e.g. from Algeria, Russia and UK. | (43) |

| Age (Mya) | Type | Description | Ref(s). |
|---|---|---|---|
| 185/190 | Minimum | The first widely recognized record of centric diatoms described from Toarcian samples of the Liassic Boll shales of Wurtenburg, Germany. The date 190 Mya refers to the calibration point from Sánchez-Baracaldo et al. (19). | (44) |
| 130 | Minimum | The microfossils of angiosperm pollen grains described from Hauterivian material of the Warlingham borehole, UK. | (45, 46) |
| 125 | Minimum | The fossil record of angiosperms, e.g. *Nymphaeaceae* or *Ceratophyllum*, goes back to the early Cretaceous about 125 Mya. | (47) |
| 110 | Minimum | The first well-preserved deposit of diatoms described from Aptian–Albian core material (115 - 110 Myr-old) recovered from the Weddell Sea, Antarctica. | (48) |

**Table S3.** Molecular clock estimates for key nodes in our analyses.

| Node | Clade | Age in million years according to the calibration set | | | Software |
|---|---|---|---|---|---|
| | | C1 | C2 | C3 | |
| I | *Gloeomargarita* + Archaeplastida | 2039.29 (2342.93-1755.29) | 2040.2 (2383.37-1762.4) | 2156.46 (2440.91-1843.44) | MrBayes IGR |
| | | 1926.77 (2045.7-1817.55) | 2132.25 (2311.24-1954.47) | 2207.3 (2370.7-2053.59) | MrBayes TK02 |
| | | 1929.82 (2172.05-1589.29) | 2044.85 (2362.47-1712.71) | 2150.93 (2416.38-1916.71) | Beast UCLNR |
| | | 2172 (2343.28-2016.44) | 2189 (2374.48-2029.18) | 2196 (2382.01-2044.88) | Phylobayes UGAM |
| | | 1937 (2016.44-1853.27) | 1947 (2047.2-1858.44) | 1986 (2088.69-1914.05) | Phylobayes CIR |
| | | 1951 (2040.66-1848.74) | 2066 (2206.72-1947.44) | 2090 (2240.55-1981.71) | Phylobayes LN |
| II | Divergence of *Glaucophyta* | 1943.82 (2232.79-1674.82) | 1938.39 (2271.99-1644.06) | 2066.61 (2335.76-1797.55) | MrBayes IGR |
| | | 1673.76 (1783.11-1569.51) | 1884.4 (2027.79-1746.16) | 1961.93 (2084.77-1849.59) | MrBayes TK02 |
| | | 1765.99 (2015.53-1528.39) | 1879.78 (2172.08-1568.27) | 1989.09 (2218.78-1785.02) | Beast UCLNR |
| | | 2057 (2237.87-1883.46) | 2075 (2263.76-1901.99) | 2082 (2271.52-1917.57) | Phylobayes UGAM |
| | | 1760 (1847.86-1662.12) | 1780 (1880.16-1692.45) | 1827 (1915.73-1768.82) | Phylobayes CIR |
| | | 1772 (1869.1-1641.83) | 1929 (2063.08-1814.06) | 1953 (2074.21-1827.03) | Phylobayes LN |
| III | *Rhodophyta* + Chloroplastida | 1865.54 (2159.6-1615.89) | 1860.63 (2199.38-1582.97) | 1991.06 (2272.93-1710.75) | MrBayes IGR |
| | | 1617.86 (1727.17-1512.74) | 1830.81 (1966.04-1703.3) | 1911.82 (2024.92-1808.12) | MrBayes TK02 |
| | | 1671.79 (1930.5-1429.17) | 1786.14 (2043.62-1445.63) | 1899.1 (2093.84-1708.03) | Beast UCLNR |
| | | 2023 (2206.7-1845.08) | 2041 (2232.54-1863.43) | 2047 (2238.14-1879.27) | Phylobayes UGAM |
| | | 1748 (1838.96-1648.34) | 1768 (1869.15-1679.89) | 1816 (1902.12-1759.08) | Phylobayes CIR |
| | | 1761 (1860.91-1625.52) | 1921 (2053.56-1806.78) | 1945 (2064.79-1816.96) | Phylobayes LN |
| IV | Crown group Chloroplastida | 1734.31 (1980.26-1494.98) | 1738.38 (2045.77-1465.15) | 1842.62 (2128.78-1555.16) | MrBayes IGR |
| | | 1333.18 (1434.37-1235.94) | 1542.71 (1636.32-1455.86) | 1613.12 (1696.34-1538.08) | MrBayes TK02 |
| | | 1467.81 (1746.76-1237.7) | 1619.35 (1860.73-1354.01) | 1704.88 (1935.74-1495.19) | Beast UCLNR |
| | | 1812 (2019.18-1599.16) | 1832 (2039.2-1628.45) | 1837 (2044.86-1637.09) | Phylobayes UGAM |
| | | 1397 (1528.58-1255.47) | 1454 (1545.17-1374.24) | 1501 (1586.33-1427.86) | Phylobayes CIR |
| | | 1403 (1582.37-1206.65) | 1672 (1802.9-1565.96) | 1685 (1791.27-1576.43) | Phylobayes LN |
| V | Crown group *Rhodophyta* | 1561.2 (1833.6-1249.82) | 1556.62 (1877.42-1264.31) | 1725.66 (1946.86-1560) | MrBayes IGR |
| | | 1243.13 (1381.37-1083.67) | 1438.21 (1590.67-1257.29) | 1589.55 (1642.73-1560) | MrBayes TK02 |
| | | 1366.18 (1669.52-1040.01) | 1345.37 (1633.12-1081.83) | 1631.66 (1761.57-1560) | Beast UCLNR |
| | | 1734 (1983.99-1436.78) | 1748 (2002.26-1454.58) | 1775 (2008.19-1579.91) | Phylobayes UGAM |
| | | 1495 (1608.57-1365.58) | 1528 (1641.91-1419.17) | 1594 (1663.33-1561.13) | Phylobayes CIR |
| | | 1525 (1660.16-1358.63) | 1726 (1868.43-1581.22) | 1753 (1859.33-1631.2) | Phylobayes LN |
| VI | Crown group *Chlorophyta* | 1573.77 (1798.44-1338.25) | 1563.82 (1854.79-1323.15) | 1684.43 (1937.8-1403.17) | MrBayes IGR |
| | | 1182.84 (1270.39-1096.79) | 1378.07 (1454.4-1306.17) | 1427.42 (1503.76-1356.86) | MrBayes TK02 |
| | | 1251.99 (1499.09-1023.75) | 1428.38 (1656.37-1201.08) | 1500.98 (1726.29-1290.89) | Beast UCLNR |
| | | 1601 (1828.34-1368.07) | 1624 (1846.25-1414.53) | 1627 (1850.09-1417.64) | Phylobayes UGAM |
| | | 1204 (1343.19-1063.81) | 1274 (1353.18-1211.39) | 1304 (1395.83-1227.31) | Phylobayes CIR |
| | | 1129 (1336.33-986.17) | 1425 (1528.42-1342.52) | 1429 (1539.14-1338.81) | Phylobayes LN |
| VII | Divergence of *Cryptophyta* | 1407.77 (1713.89-1129.5) | 1396.14 (1710.32-1085.27) | 1557.89 (1820.99-1324.36) | MrBayes IGR |
| | | 1140.74 (1304.45-963.74) | 1315.8 (1511.17-1084.34) | 1490.19 (1572.41-1413.96) | MrBayes TK02 |
| | | 1223.85 (1585.08-886.72) | 1183.98 (1535.52-914.64) | 1417.57 (1649.53-1132.57) | Beast UCLNR |
| | | 1578 (1870.16-1232.57) | 1593 (1888.23-1247.86) | 1616 (1893.52-1296.78) | Phylobayes UGAM |
| | | 1446 (1564.29-1306.6) | 1477 (1595.15-1356.91) | 1547 (1619.29-1500.88) | Phylobayes CIR |
| | | 1475 (1618.11-1299.68) | 1672 (1824.28-1508.65) | 1701 (1809.17-1567.2) | Phylobayes LN |
| VIII | Crown group *Streptophyta* | 1066.49 (1386.76-762.54) | 1111.18 (1407.31-806.47) | 1207.99 (1517.29-929.74) | MrBayes IGR |
| | | 893.29 (954.49-832.3) | 1017.32 (1088.87-946.38) | 1039.51 (1115.72-964.75) | MrBayes TK02 |
| | | 934.62 (1252.09-628.56) | 977.08 (1285.09-653.5) | 1038.51 (1399.84-676.33) | Beast UCLNR |
| | | 1100 (1513.05-721.91) | 1109 (1521.87-734.58) | 1114 (1525.37-738.7) | Phylobayes UGAM |

| Node | Clade | Age in million years according to the calibration set | | | Software |
|---|---|---|---|---|---|
| | | C1 | C2 | C3 | |
| | | 647 (703.68-607.41) | 670 (722.44-629.42) | 669 (725.81-628) | Phylobayes CIR |
| | | 651 (731-597.1) | 727 (862.59-640) | 725 (856.23-641.43) | Phylobayes LN |
| IX | Divergence of *Chlorarachnea* | 1278.52 (1496.93-1065.85) | 1261.96 (1500.83-1011.07) | 1361.38 (1596.18-1123.31) | MrBayes IGR |
| | | 786.3 (872.12-694.81) | 1013.77 (1049.55-983.86) | 1029.02 (1078.44-989.03) | MrBayes TK02 |
| | | 865.33 (1043.84-672.93) | 1076.52 (1219.55-960.21) | 1106.88 (1272.62-969.12) | Beast UCLNR |
| | | 1132 (1377.85-900.72) | 1164 (1389.28-990.69) | 1164 (1389.13-989.93) | Phylobayes UGAM |
| | | 914 (1048.77-780.98) | 991 (1058.92-954.61) | 1013 (1104.08-957.52) | Phylobayes CIR |
| | | 741 (928-649) | 995 (1059.57-954.78) | 999 (1073.15-955.83) | Phylobayes LN |
| X | Cyanobium + *Paulinella* | 450.67 (658.98-283.24) | 416.01 (594.64-218.22) | 442.6 (667.69-248.5) | MrBayes IGR |
| | | 425.31 (592.21-291.33) | 442.37 (611.57-286.99) | 463.08 (617.13-306.18) | MrBayes TK02 |
| | | 424.81 (692.39-190.55) | 434.21 (715.68-189.82) | 481.24 (772.09-234.4) | Beast UCLNR |
| | | 511 (1109.22-203.27) | 512 (1102.67-207.69) | 516 (1115.18-207.52) | Phylobayes UGAM |
| | | 156 (203.16-116.82) | 159 (208.12-119.13) | 158 (208.87-117.32) | Phylobayes CIR |
| | | 240 (308.37-166.53) | 259 (335.02-177.78) | 266 (344.18-183.87) | Phylobayes LN |
| XI | Divergence of *Paulinella* | 298.39 (456.31-157.48) | 278.82 (436-140.53) | 291.97 (440.69-162.8) | MrBayes IGR |
| | | 259.74 (417.13-145.19) | 263.91 (433.97-136.33) | 279.23 (419.01-154.71) | MrBayes TK02 |
| | | 231.31 (386.95-107.74) | 239.99 (419.71-85.72) | 275.32 (505.49-99.24) | Beast UCLNR |
| | | 264 (656.37-95.44) | 264 (645.15-96.98) | 266 (652.95-96.6) | Phylobayes UGAM |
| | | 66 (92.45-46.33) | 68 (94.82-47.36) | 67 (94.31-46.18) | Phylobayes CIR |
| | | 106 (149.5-63.99) | 114 (163.42-70.23) | 118 (170.22-71.86) | Phylobayes LN |

**Table S4.** List of 30 conserved plastid-encoded proteins and query species used for homologous search by PSI-BLAST.

| Gene | Protein | NCBI accession number | | | | |
|---|---|---|---|---|---|---|
| | | ***Arabidopsis thaliana*** | ***Cyanobium sp.* NIES-981** | ***Cyanophora paradoxa*** | ***Galdieria sulphuraria*** | ***Gloeomargarita lithophora* Alchichica-D10** |
| *atpA* | ATP synthase subunit alpha | ANW47776.1 | SBO44543.1 | NP_043222.1 | AIG92565.1 | APB33236.1 |
| *atpB* | ATP synthase subunit beta | NP_051066.1 | WP_087068740.1 | NP_043241.1 | AIG92549.1 | APB34992.1 |
| *atpH* | ATP synthase subunit c | ANW47777.1 | SBO44542.1 | NP_043226.1 | AIG92561.1 | WP_071453852.1 |
| *ccsA* | Cytochrome c biogenesis protein | NP_051108.1 | SBO42325.1 | NP_043267.1 | P31564.1 | APB32776.1 |
| *petA* | Component of the cytochrome b6-f complex | ANW47803.1 | SBO42987.1 | NP_043244.1 | AIG92543.1 | APB32505.1 |
| *petB* | Component of the cytochrome b6-f complex | NP_051088.1 | SBO43464.1 | NP_043175.1 | AIG92473.1 | APB34594.1 |
| *petD* | Component of the cytochrome b6-f complex | NP_051089.1 | SBO43463.1 | NP_043174.1 | AIG92474.1 | APB34593.1 |
| *psaA* | Photosystem I P700 chlorophyll a apoprotein A1 | NP_051059.1 | SBO43593.1 | AAA81181.1 | AGZ04878.1 | APB33122.1 |
| *psaC* | Photosystem I iron-sulfur center | ANW47840.1 | SBO43969.1 | AAA81301.1 | AIG92625.1 | WP_071454602.1 |
| *psbA* | Photosystem II protein D1 | CAA56270.1 | SBO44640.1 | NP_043238.1 | AIG92634.1 | APB33505.1 |
| *psbB* | Photosystem II CP47 reaction center protein | ANW47816.1 | SBO43481.1 | AAA81198.1 | AIG92640.1 | APB33253.1 |
| *psbC* | Photosystem II CP43 reaction center protein | ANW47786.1 | SBO44375.1 | NP_043248.3 | AIG92464.1 | APB33217.1 |
| *psbD* | Photosystem II D2 protein | ANW47785.1 | SBO44818.1 | NP_043247.1 | AIG92463.1 | APB33216.1 |
| *psbE* | Cytochrome b559 subunit alpha | NP_051076.1 | SBO43722.1 | NP_043178.1 | AIG92575.1 | APB32556.1 |
| *psbK* | Photosystem II reaction center protein K | ANW47774.1 | SBO43700.1 | AAA81280.1 | AIG92530.1 | WP_071454683.1 |
| *rbcL* | Ribulose bisphosphate carboxylase large chain | AAB68400.1 | SBO42577.1 | NP_043240.1 | AGZ04769.1 | APB32635.1 |
| *rpl14* | 50S ribosomal protein L14, chloroplastic | ANW47826.1 | WP_087068059.1 | AAA63624.1 | AIG92509.1 | APB34322.1 |
| *rpL20* | 50S ribosomal protein L20, chloroplastic | NP_051082.1 | SBO44063.1 | NP_043162.1 | AIG92539.1 | APB32669.1 |
| *rpoA* | DNA-directed RNA polymerase subunit alpha | NP_051090.1 | SBO43574.1 | NP_043256.1 | AIG92499.1 | APB34409.1 |
| *rpoB* | DNA-directed RNA polymerase subunit beta | CAA74024.1 | SBO44625.1 | NP_043230.1 | AIG92555.1 | APB34630.1 |
| *rpoC1* | DNA-directed RNA polymerase subunit beta' | NP_051050.1 | WP_087068805.1 | NP_043229.1 | AIG92556.1 | WP_071455042.1 |
| *rpoC2* | DNA-directed RNA polymerase subunit beta'' | NP_051049.1 | WP_087068804.1 | NP_043228.1 | AIG92557.1 | APB34628.1 |
| *S11* | 30S ribosomal protein S11, chloroplastic | NP_051091.1 | SBO43573.1 | NP_043257.1 | AIG92500.1 | APB34410.1 |
| *S12* | 30S ribosomal protein S12, chloroplastic | ALE59953.2 | SBO43601.1 | NP_043209.1 | AIG92495.1 | APB33533.1 |
| *S19* | 30S ribosomal protein S19, chloroplastic | NP_051098.1 | SBO43555.1 | NP_043198.1 | AIG92514.1 | APB34316.1 |
| *S2* | 30S ribosomal protein S2, chloroplastic | NP_051048.1 | SBO42340.1 | NP_043227.1 | AIG92558.1 | APB35091.1 |
| *S3* | 30S ribosomal protein S3, chloroplastic | NP_051096.1 | WP_087069299.1 | NP_043196.1 | AIG92512.1 | APB34318.1 |
| *S4* | 30S ribosomal protein S4, chloroplastic | ANW47792.1 | WP_087067709.1 | NP_043212.1 | AIG92603.1 | APB34495.1 |
| *S7* | 30S ribosomal protein S7, chloroplastic | NP_051118.1 | SBO43602.1 | NP_043208.1 | AIG92494.1 | APB33532.1 |
| *ycf3* | Photosystem I assembly protein | BAA84386.1 | SBO44825.1 | AAA81185.1 | Q08814.1 | WP_071453871.1 |

**Table S5.** Partitions and substitution models proposed by ModelFinder (49) for IQ-TREE (50).

| Model | Partition no. | Partition range |
|---|---|---|
| LGF | Subset1 | 5116-5236, 4484-4564 1-500 8530-8658 |
| cpREV | Subset2 | 501-960 8659-8779 |
| mtZOAF | Subset3 | 961-1039 2824-3175 2727-2823 4133-4483 1829-1987 1988-2726 3684-4132 |
| cpREVF | Subset4 | 4565-4626 1040-1294 1295-1606 |
| LG | Subset5 | 1607-1828 4627-5115 3176-3683 |
| cpREV | Subset6 | 5237-5350 9095-9302 |
| WAGF | Subset7_Subset12 | 5351-5649 7228-8529 |
| cpREV | Subset8 | 5650-6623 6624-7227 8871-9094 9303-9501 |
| cpREV | Subset9 | 8780-8870 9502-9656, 9657-9823 |

**Table S6.** Partitions and substitution models proposed by Partition finder (51) for RAxML (52).

| Model | Partition no. | Partition range |
|---|---|---|
| LGF | Subset1 | 5116-5236 4484-4564 1-500 |
| CPREVF | Subset2 | 501-960 |
| LGF | Subset3 | 961-1039 2824-3175 |
| CPREVF | Subset4 | 4565-4626 1040-1294 |
| LGF | Subset5 | 1295-1606 |
| LG | Subset6 | 1607-1828 4627-5115 3176-3683 |
| LGF | Subset7 | 2727-2823 4133-4483 1829-1987 1988-2726 3684-4132 |
| CPREV | Subset8 | 5237-5350 9095-9302 |
| CPREV | Subset9 | 5351-5649 |
| CPREV | Subset10 | 5650-6623 |
| CPREV | Subset11 | 6624-7227 |
| WAGF | Subset12 | 7228-8529 |
| JTT | Subset13 | 8530-8658 |
| CPREV | Subset14 | 8659-8779 |
| LG | Subset15 | 8780-8870 |
| CPREV | Subset16 | 8871-9094 |
| LG | Subset17 | 9303-9501 |
| CPREV | Subset18 | 9502-9656 |
| CPREV | Subset19 | 9657-9823 |

**Table S7.** Partitions and substitution models proposed by Partition finder (51) for MrBayes (53).

| Model | Partition no. | Partition range |
|---|---|---|
| WAG | Subset1 | 5116-5236 4484-4564 1-500 |
| CPREV | Subset2 | 501-960 |
| WAG | Subset3 | 961-1039 2824-3175 |
| CPREV | Subset4 | 4565-4626 1040-1294 |
| WAG | Subset5 | 1295-1606 |
| WAG | Subset6 | 1607-1828 4627-5115 3176-3683 |
| WAG | Subset7 | 2727-2823 4133-4483 1829-1987 1988-2726 3684-4132 |
| CPREV | Subset8 | 5237-5350 9095-9302 |
| CPREV | Subset9 | 5351-5649 |
| CPREV | Subset10 | 5650-6623 |
| CPREV | Subset11 | 6624-7227 |
| WAG | Subset12 | 7228-8529 |
| JTT | Subset13 | 8530-8658 |
| CPREV | Subset14 | 8659-8779 |
| WAG | Subset15 | 8780-8870 |
| CPREV | Subset16 | 8871-9094 |
| WAG | Subset17 | 9303-9501 |
| CPREV | Subset18 | 9502-9656 |
| CPREV | Subset19 | 9657-9823 |

**Table S8.** Partitions and substitution models proposed by Partition finder (51) for Beast (54).

| Model | Partition no. | Partition range |
|---|---|---|
| LG+I+G | Subset1 | 1-500 |
| LG+I+G | Subset2 | 501-960 |
| LG+I+G | Subset3 | 4133-4483 961-1039 2824-3175 |
| CPREV+G | Subset4 | 1040-1294 |
| LG+I+G | Subset5 | 1295-1606 |
| LG+I+G | Subset6 | 1607-1828 1988-2726 |
| LG+I+G | Subset7 | 3684-4132 1829-1987 |
| LG+I+G | Subset8 | 2727-2823 |
| LG+I+G | Subset9 | 4627-5115 3176-3683 4484-4564 |
| CPREV+I+G | Subset10 | 5237-5350 4565-4626 |
| LG+I+G | Subset11 | 5116-5236 |
| CPREV+I+G | Subset12 | 5351-5649 |
| CPREV+I+G | Subset13 | 5650-6623 9095-9302 |
| CPREV+I+G | Subset14 | 6624-7227 |
| CPREV+I+G | Subset15 | 7228-8529 |
| JTT+I+G | Subset16 | 8530-8658 |
| CPREV+G | Subset17 | 8659-8779 |
| LG+I+G | Subset18 | 8780-8870 |
| CPREV+I+G | Subset19 | 8871-9094 |
| LG+I+G | Subset20 | 9303-9501 |
| CPREV+I+G | Subset21 | 9502-9656 |
| CPREV+G | Subset22 | 9657-9823 |

**Table S9.** Climatic and atmospheric parameters used to correlate with the diversification rate.

| Parameter | Period [Mya] | Reference |
|---|---|---|
| Atmospheric $CO_2$ | 4000-750; 750-0 | (55, 56) |
| Atmospheric $CO_2$ (COPSE model) | 750-0 | (55) |
| Atmospheric $CO_2$ (GEOCARBSULF model) | 570-0 | (55) |
| Atmospheric $CO_2$ | 570-0 | (57) |
| Atmospheric $CO_2$ | 550-0 | (58) |
| Atmospheric $CO_2$ | 420-0 | (59) |
| Atmospheric $CO_2$ | 414-0 | (57) |
| Atmospheric $CO_2$ | 405-0 | (60) |
| Atmospheric $O_2$ | 2900-0 | (61, 62) |
| Atmospheric $O_2$ | 2900-0 | (62) |
| Atmospheric $O_2$ | 2800-0 | (63) |
| Atmospheric $O_2$ | 2610-0 | (64) |
| Atmospheric $O_2$ | 630-0 | (64) |
| Atmospheric $O_2$ | 570-0 | (57) |
| Atmospheric $O_2$ | 550-0 | (58) |
| Oceanic temperature | 3500-520; 520-0 | (65, 66) |
| Global temperature | 550-0 | (58) |
| Global average temperature | 540-0 | (67) |
| Oceanic temperature | 520-0 | (65) |
| Mean sea surface temperature | 498-0 | (68) |
| The area of Large Igneous Province | 2865-0 | (69, 70) |

**Table S10.** Main glaciations for which the diversification rate was calculated.

| Name | Period [Mya] | Reference |
|---|---|---|
| Huronian (Makganyene) | 2450-2220 | (71) |
| Sturtian | 717.5-658.5 | (72) |
| Marinoan | 649.9-634.7 | (72) |
| Andean-Saharan (Hirnantian and Late Ordovician glaciation) | 456-428 | (73) |
| Karoo | 380-345 | (73) |
| Permian | 335-256 | (73) |

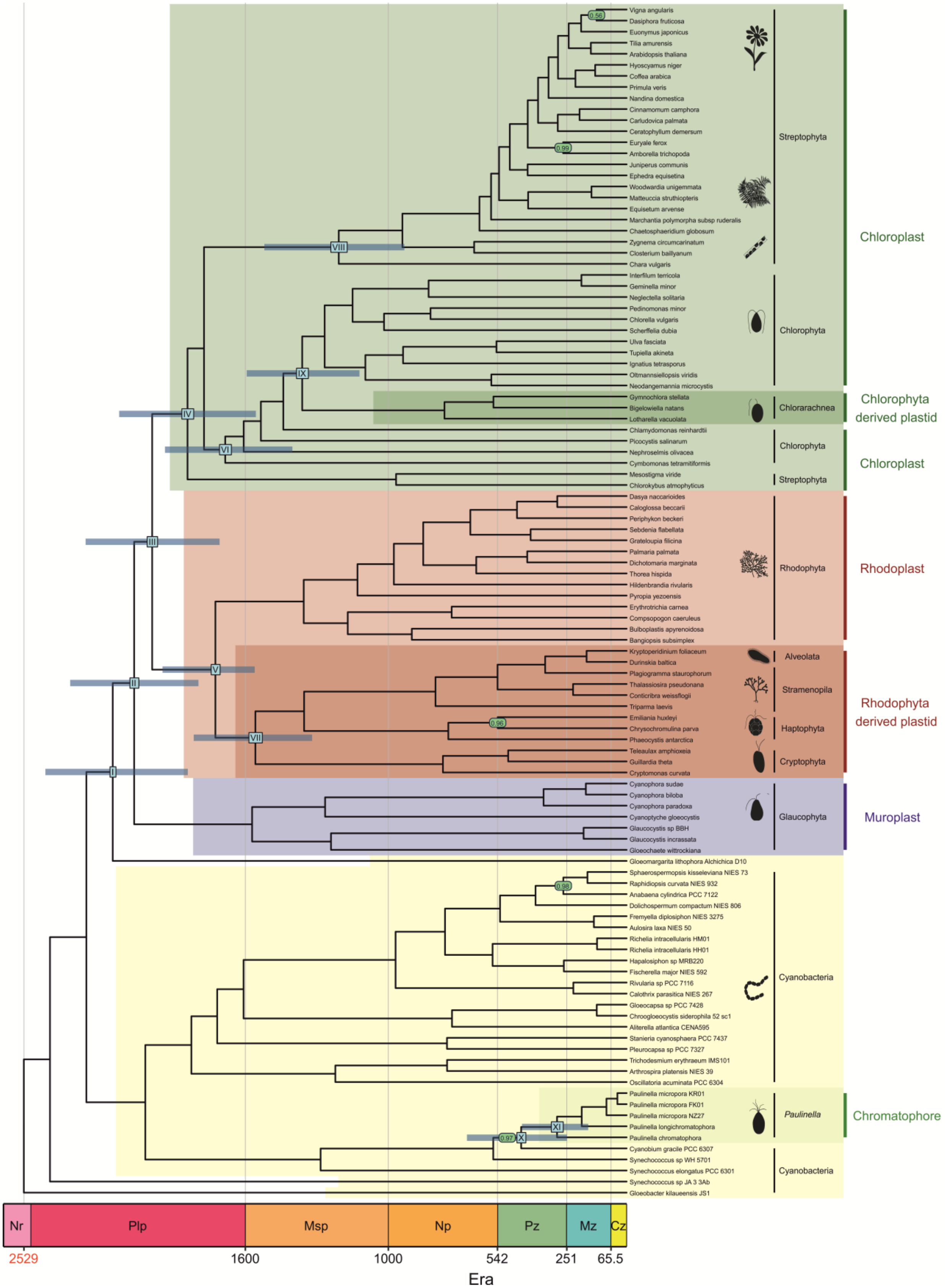


**Figure S1.** Time-calibrated phylogeny of photosynthetic organelles and cyanobacteria. The tree was inferred with MrBayes under IGR model and calibrated with C3 (Table 1). The roman numerals in blue rectangles mark key evolutionary events for plastids discussed in the article. At these nodes, there are blue bars representing 95% credibility intervals of the node age. The nodes supported with posterior probability lower than one are indicated in green circles.

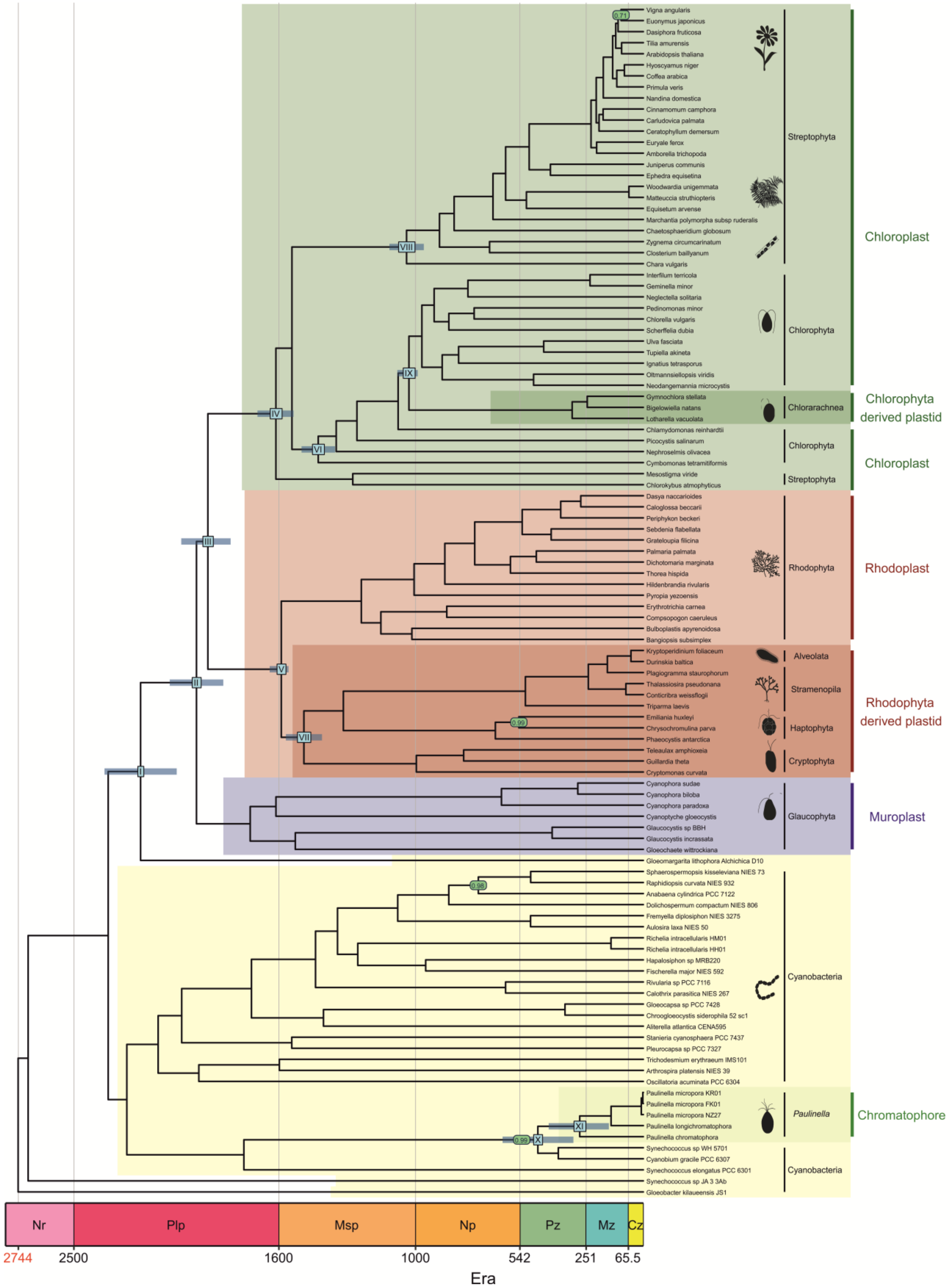


**Figure S2.** Time-calibrated phylogeny of photosynthetic organelles and cyanobacteria. The tree was inferred with MrBayes under TK02 model and calibrated with C3 (Table 1). Other description as in Fig. S1.

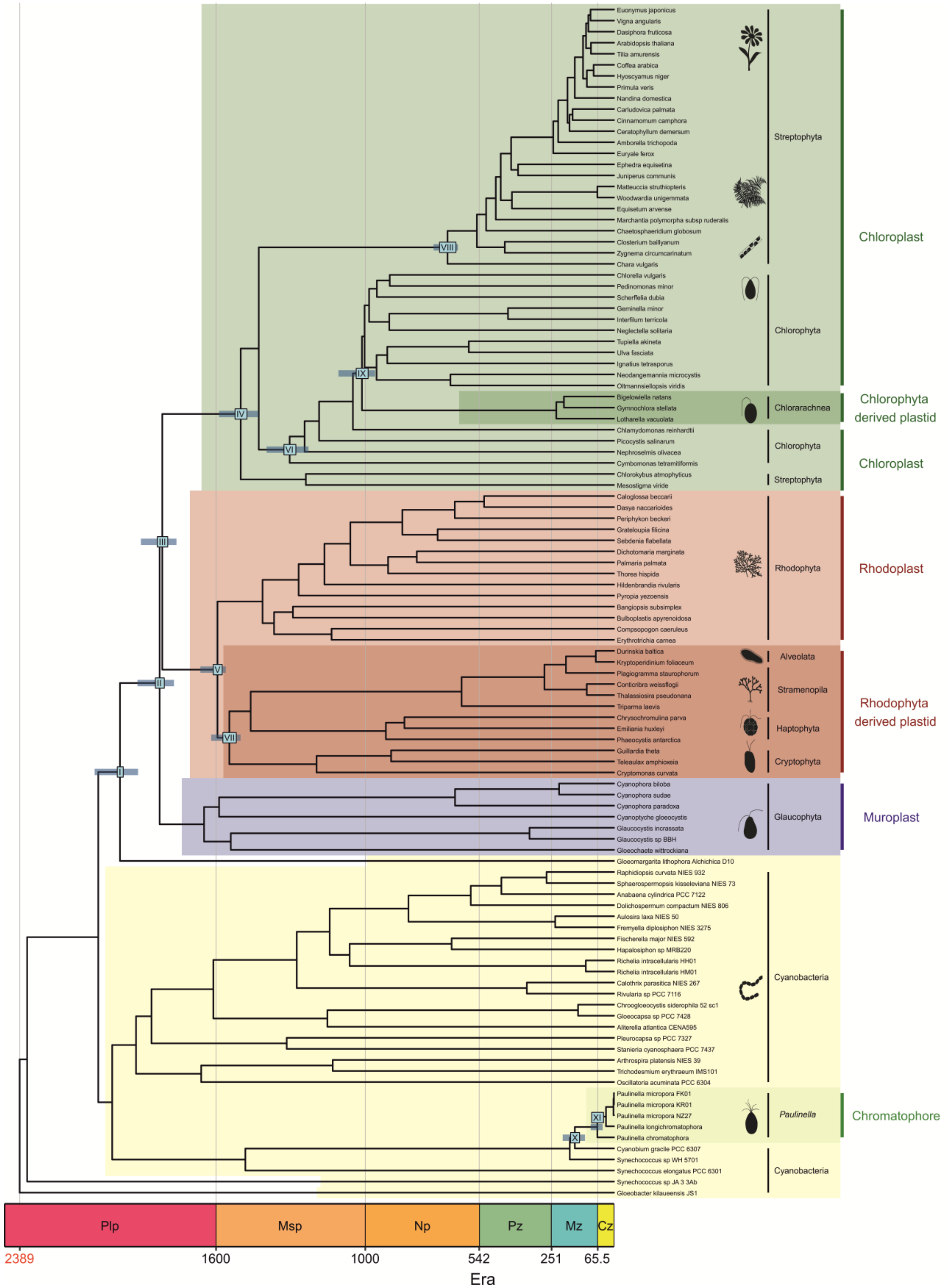


**Figure S3.** Time-calibrated phylogeny of photosynthetic organelles and cyanobacteria. The divergence times were inferred with Phylobayes under CIR model and calibrated with C3 (Table 1). The tree topology was reconstructed in IQ-TREE. Other description as in Fig. S1.

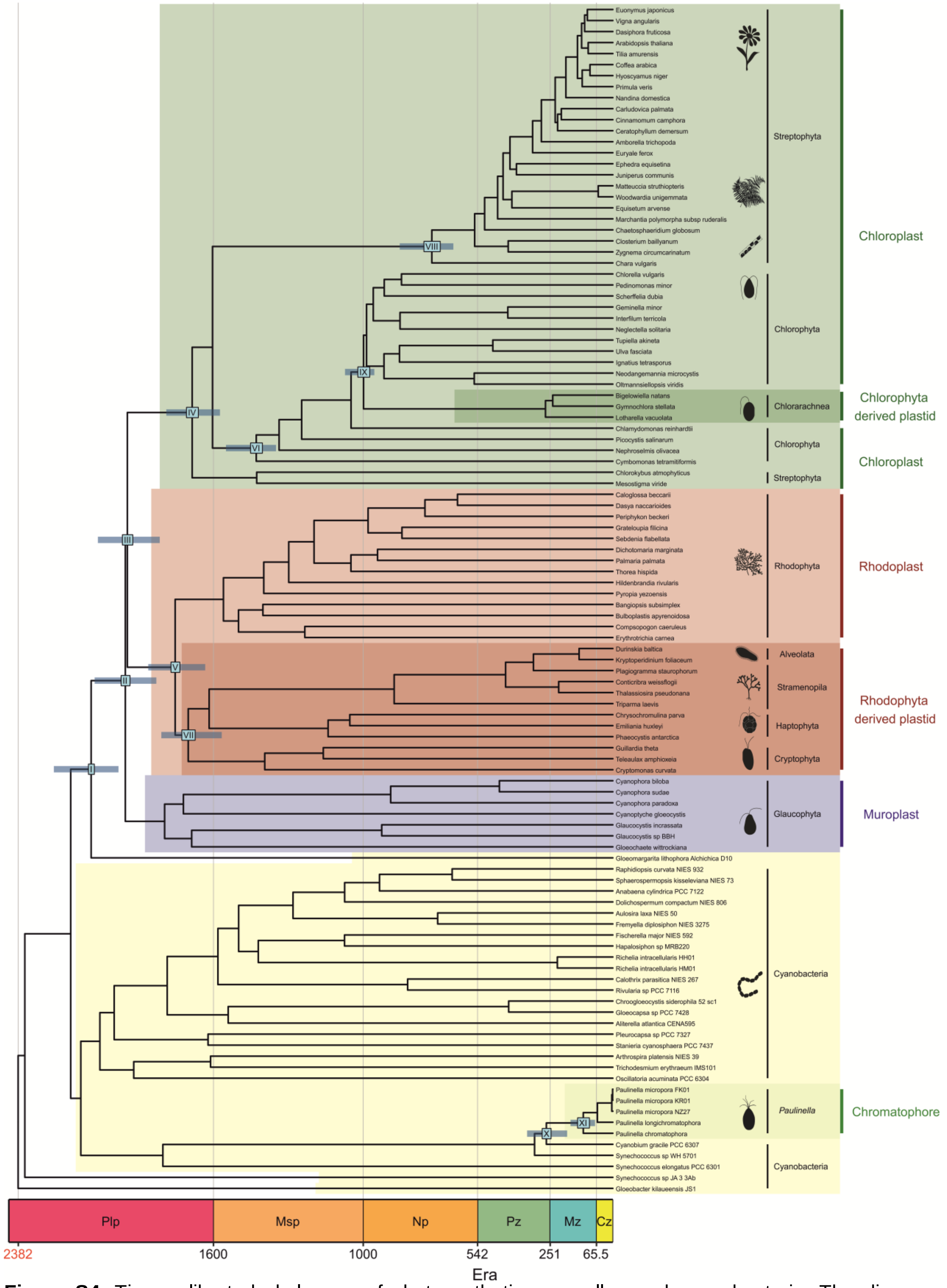


**Figure S4.** Time-calibrated phylogeny of photosynthetic organelles and cyanobacteria. The divergence times were inferred with Phylobayes under LN model and calibrated with C3 (Table 1). The tree topology was reconstructed in IQ-TREE. Other description as in Fig. S1.

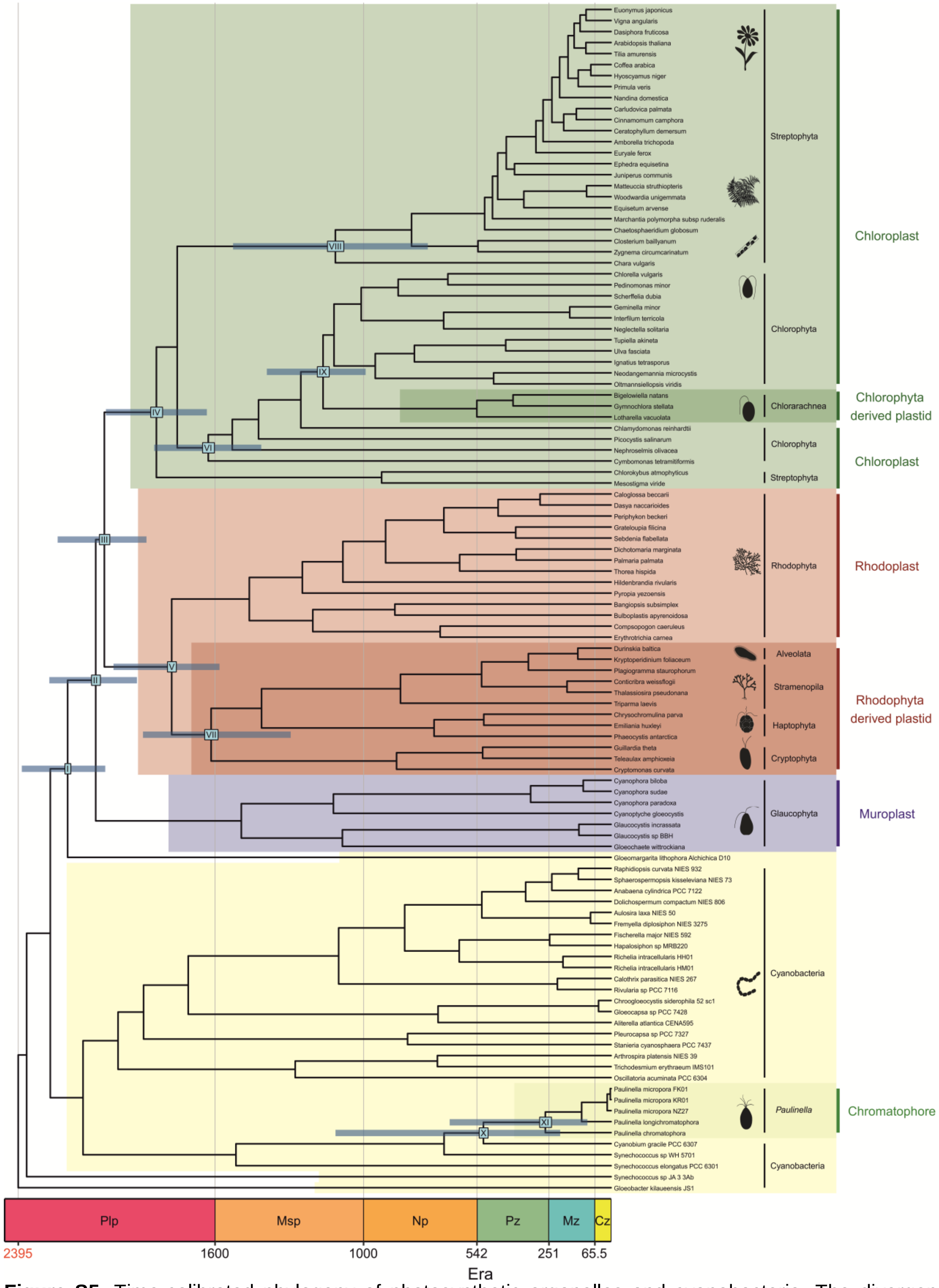


**Figure S5.** Time-calibrated phylogeny of photosynthetic organelles and cyanobacteria. The divergence times were inferred with Phylobayes under UGAM model and calibrated with C3 (Table 1). The tree topology was reconstructed in IQ-TREE. Other description as in Fig. S1.

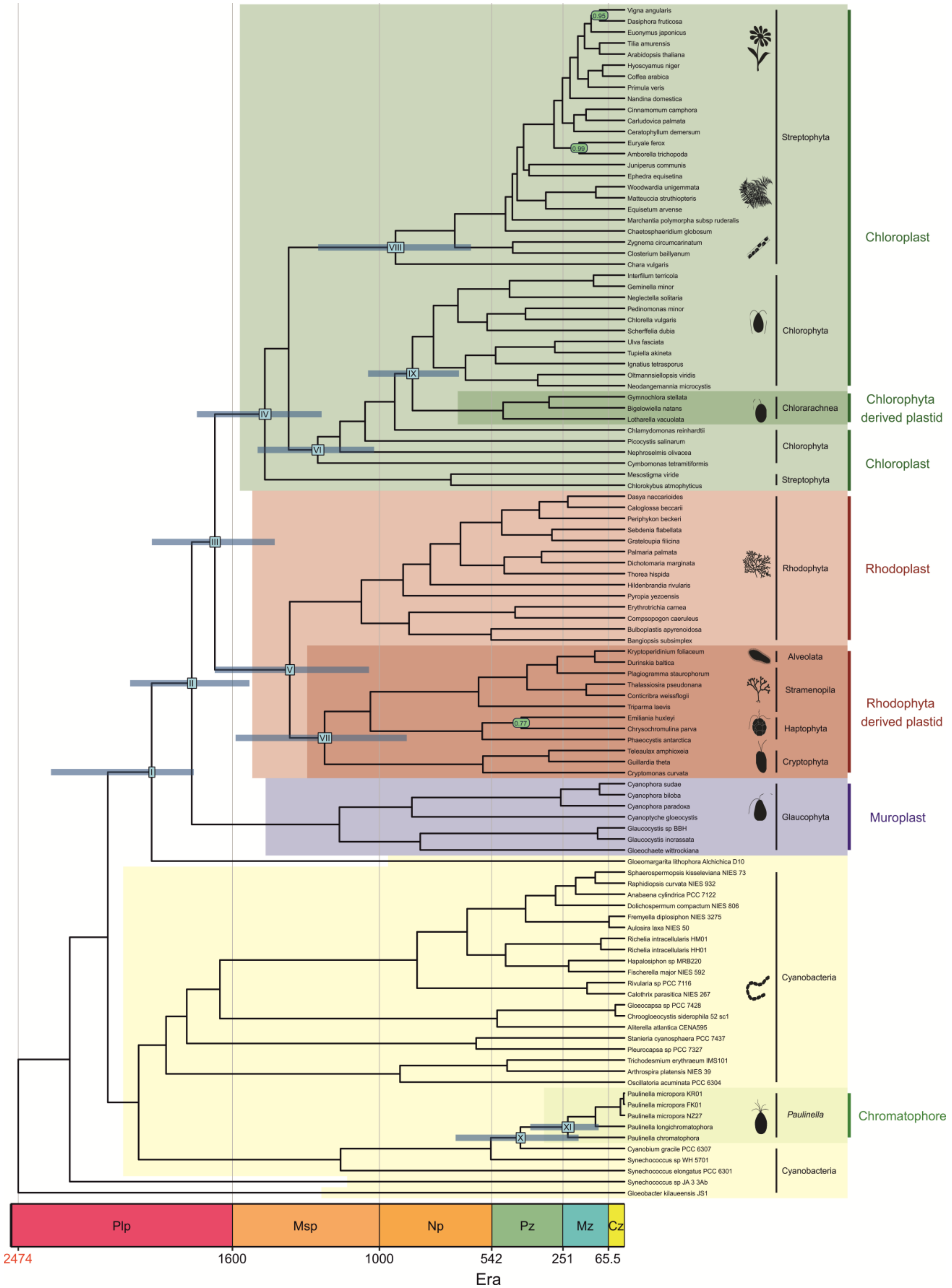


**Figure S6.** Time-calibrated phylogeny of photosynthetic organelles and cyanobacteria. The tree was inferred with Beast under UCLNR model and calibrated with C1 (Table 1). Other description as in Fig. S1.

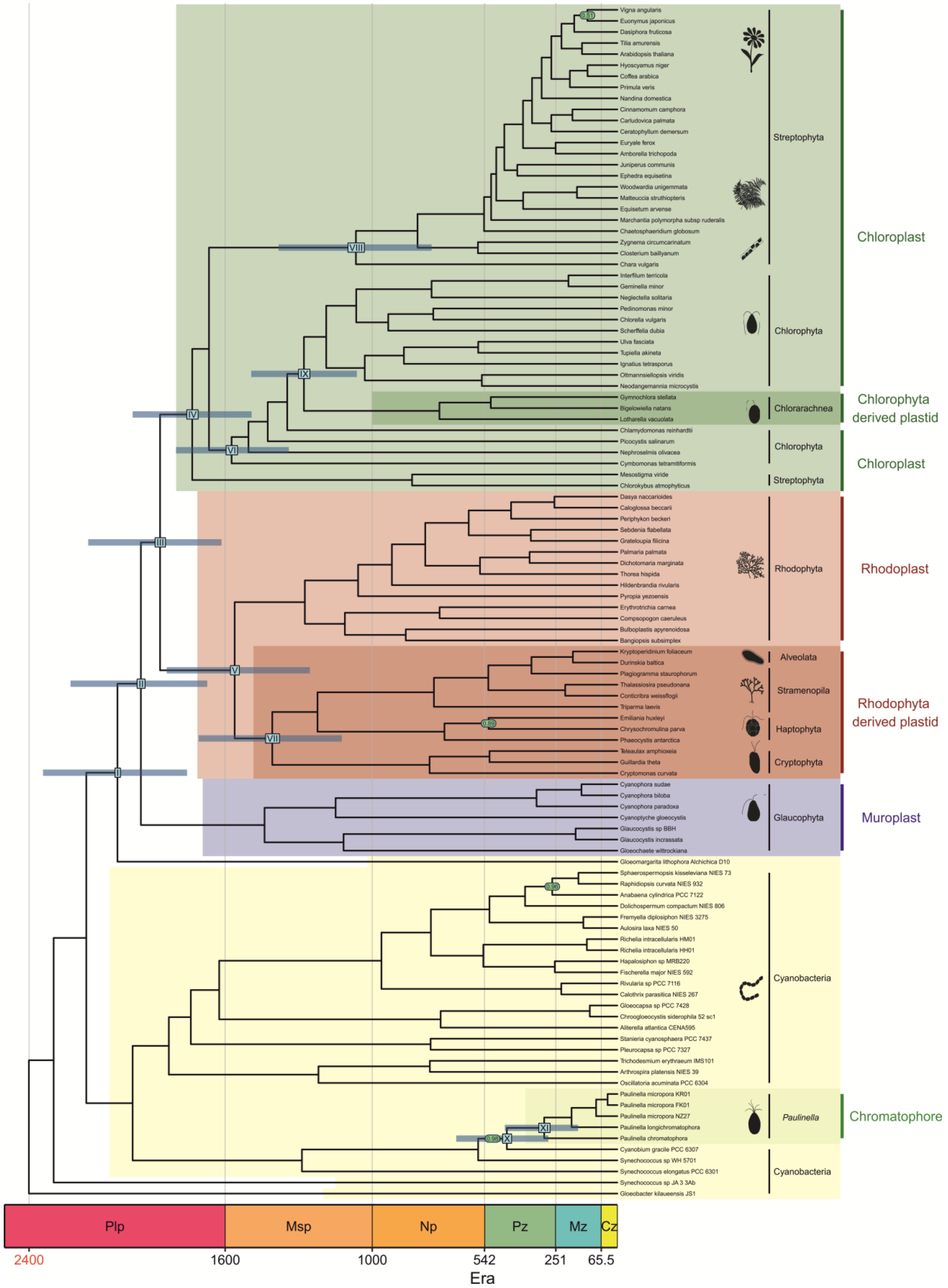


**Figure S7.** Time-calibrated phylogeny of photosynthetic organelles and cyanobacteria. The tree was inferred with MrBayes under IGR model and calibrated with C1 (Table 1). Other description as in Fig. S1.

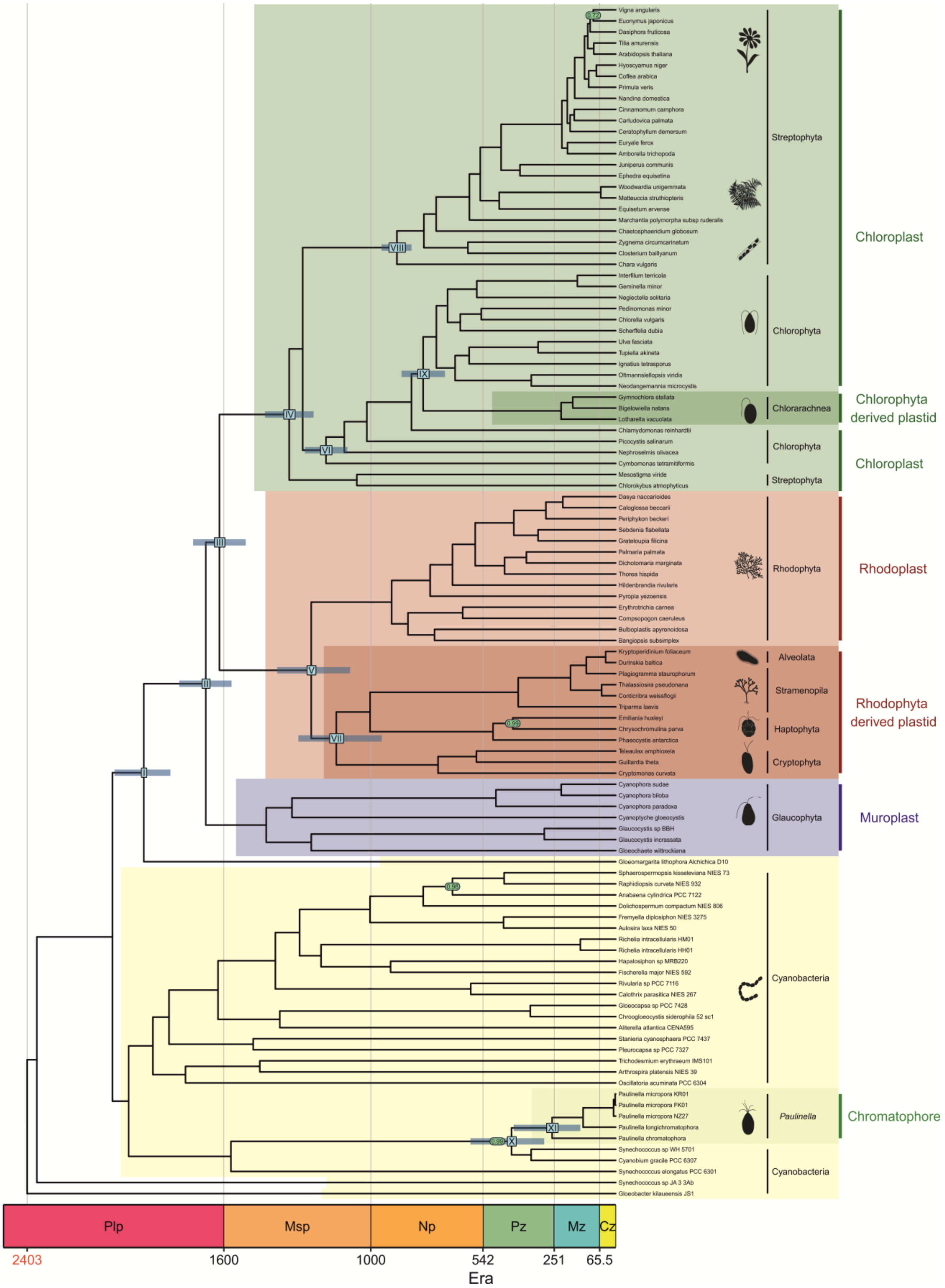


**Figure S8.** Time-calibrated phylogeny of photosynthetic organelles and cyanobacteria. The tree was inferred with MrBayes under TK02 model and calibrated with C1 (Table 1). Other description as in Fig. S1.

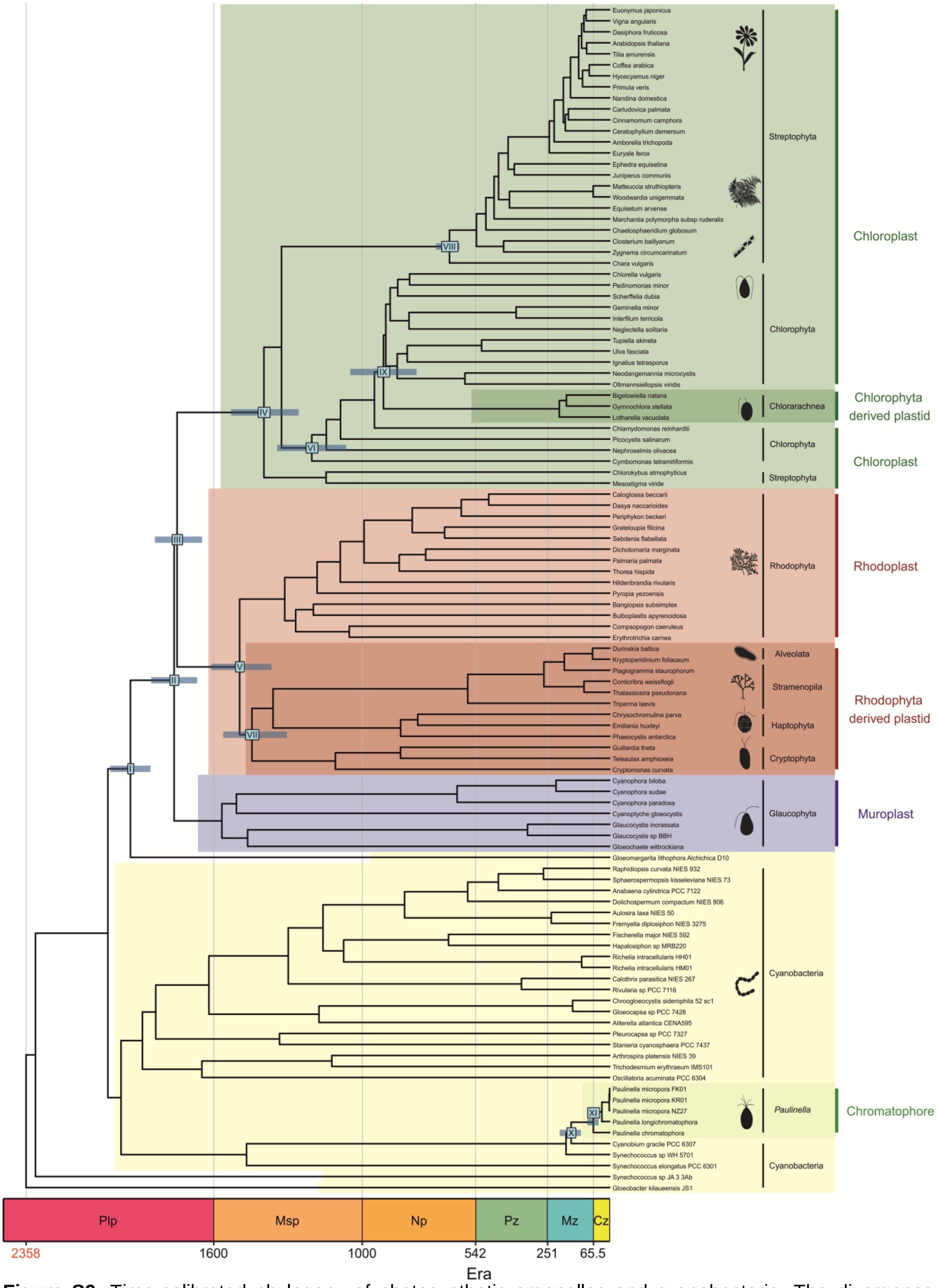


**Figure S9.** Time-calibrated phylogeny of photosynthetic organelles and cyanobacteria. The divergence times were inferred with Phylobayes under CIR model and calibrated with C1 (Table 1). The tree topology was reconstructed in IQ-TREE. Other description as in Fig. S1.

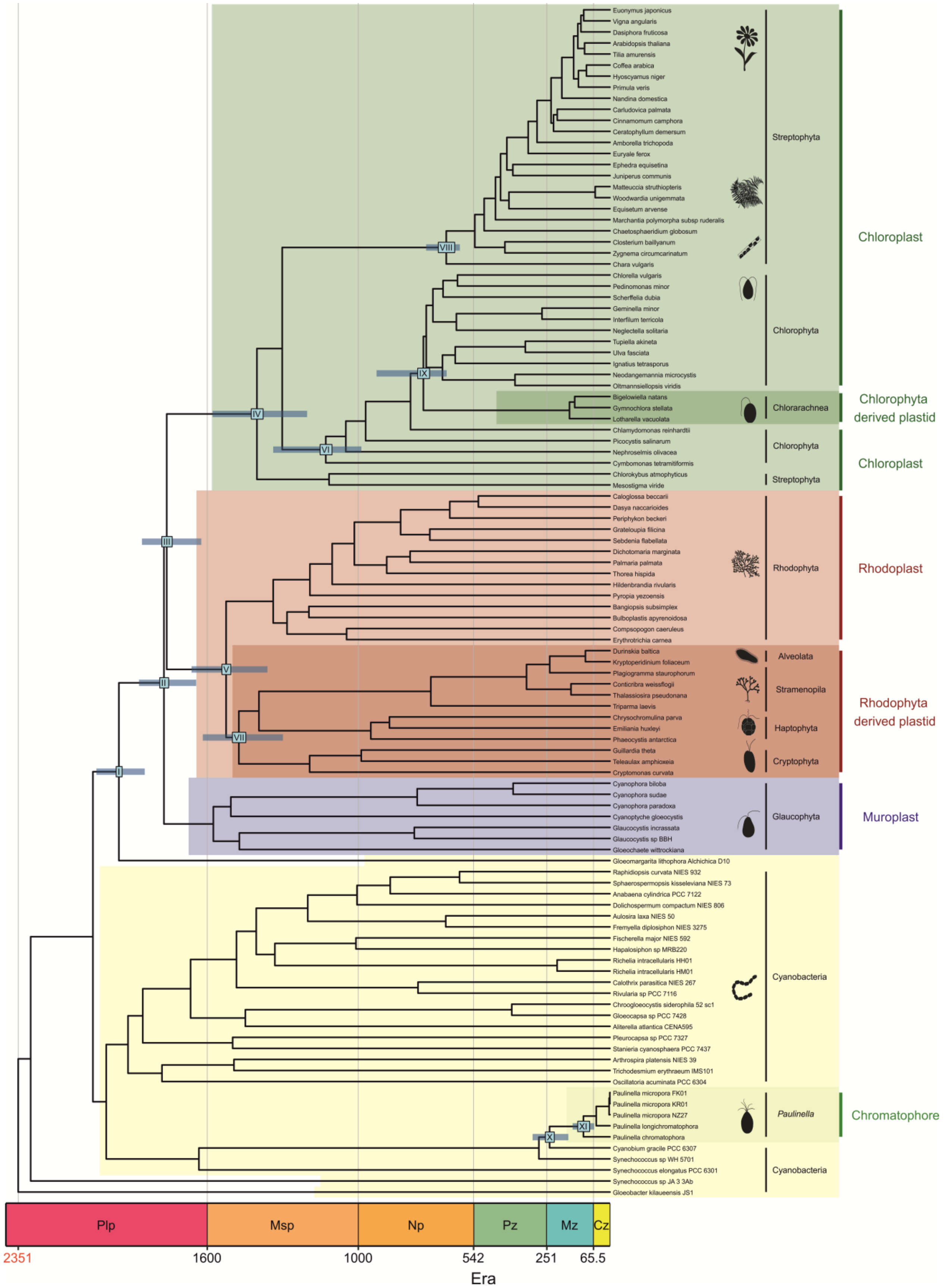


**Figure S10.** Time-calibrated phylogeny of photosynthetic organelles and cyanobacteria. The divergence times were inferred with Phylobayes under LN model and calibrated with C1 (Table 1). The tree topology was reconstructed in IQ-TREE. Other description as in Fig. S1.

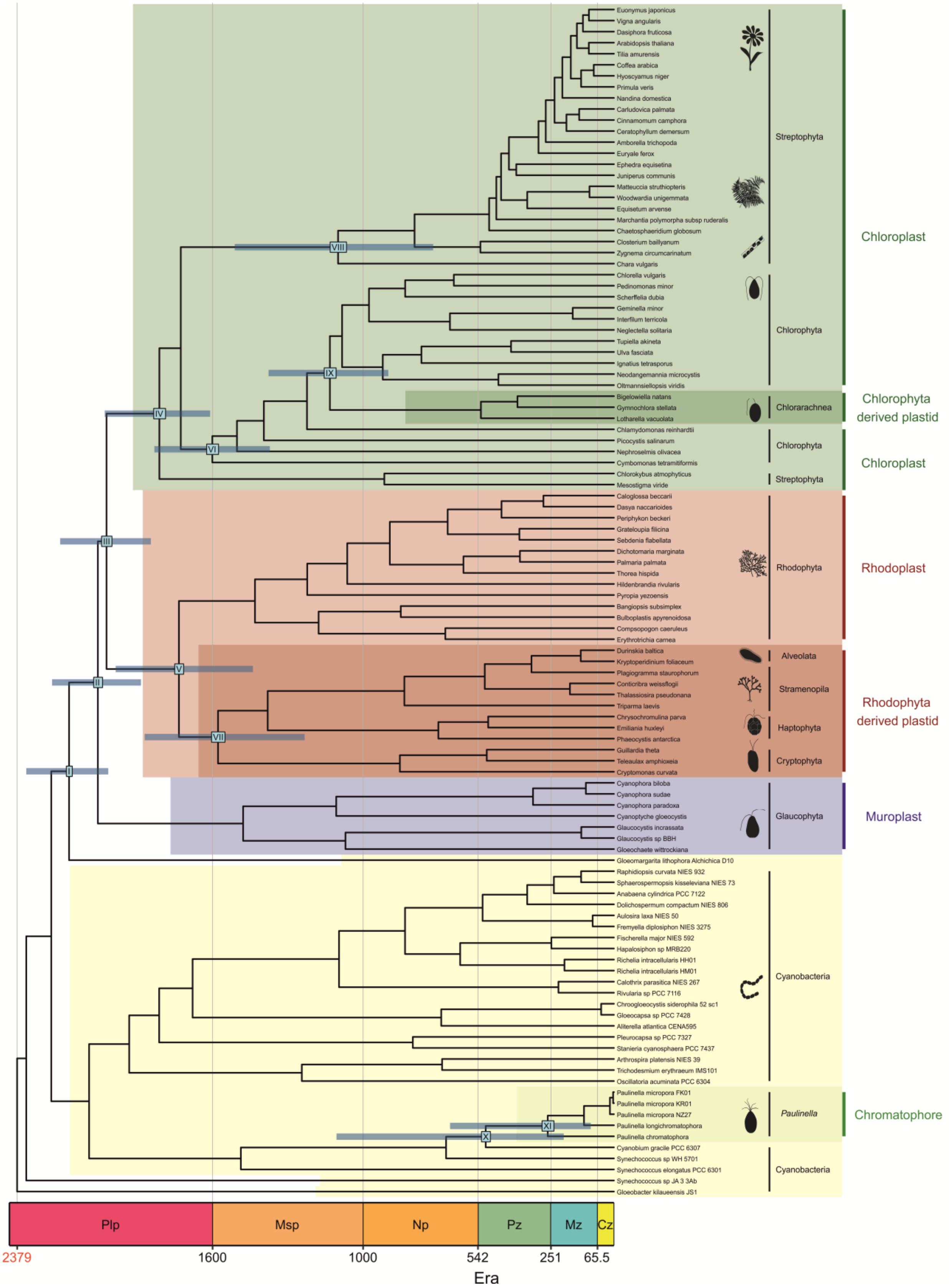


**Figure S11.** Time-calibrated phylogeny of photosynthetic organelles and cyanobacteria. The divergence times were inferred with Phylobayes under UGAM model and calibrated with C1 (Table 1). The tree topology was reconstructed in IQ-TREE. Other description as in Fig. S1.

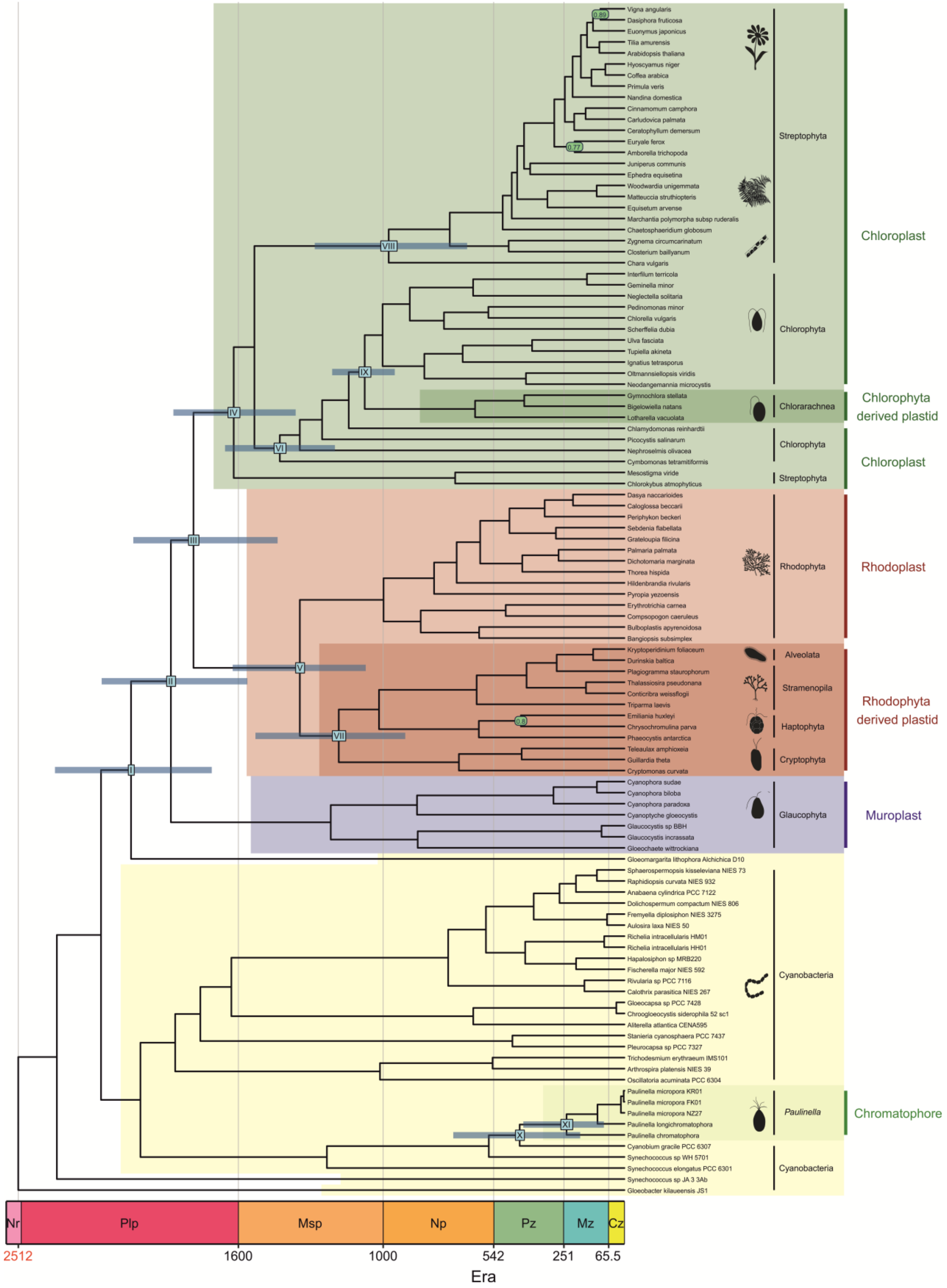


**Figure S12.** Time-calibrated phylogeny of photosynthetic organelles and cyanobacteria. The tree was inferred with Beast under UCLNR model and calibrated with C2 (Table 1). Other description as in Fig. S1.

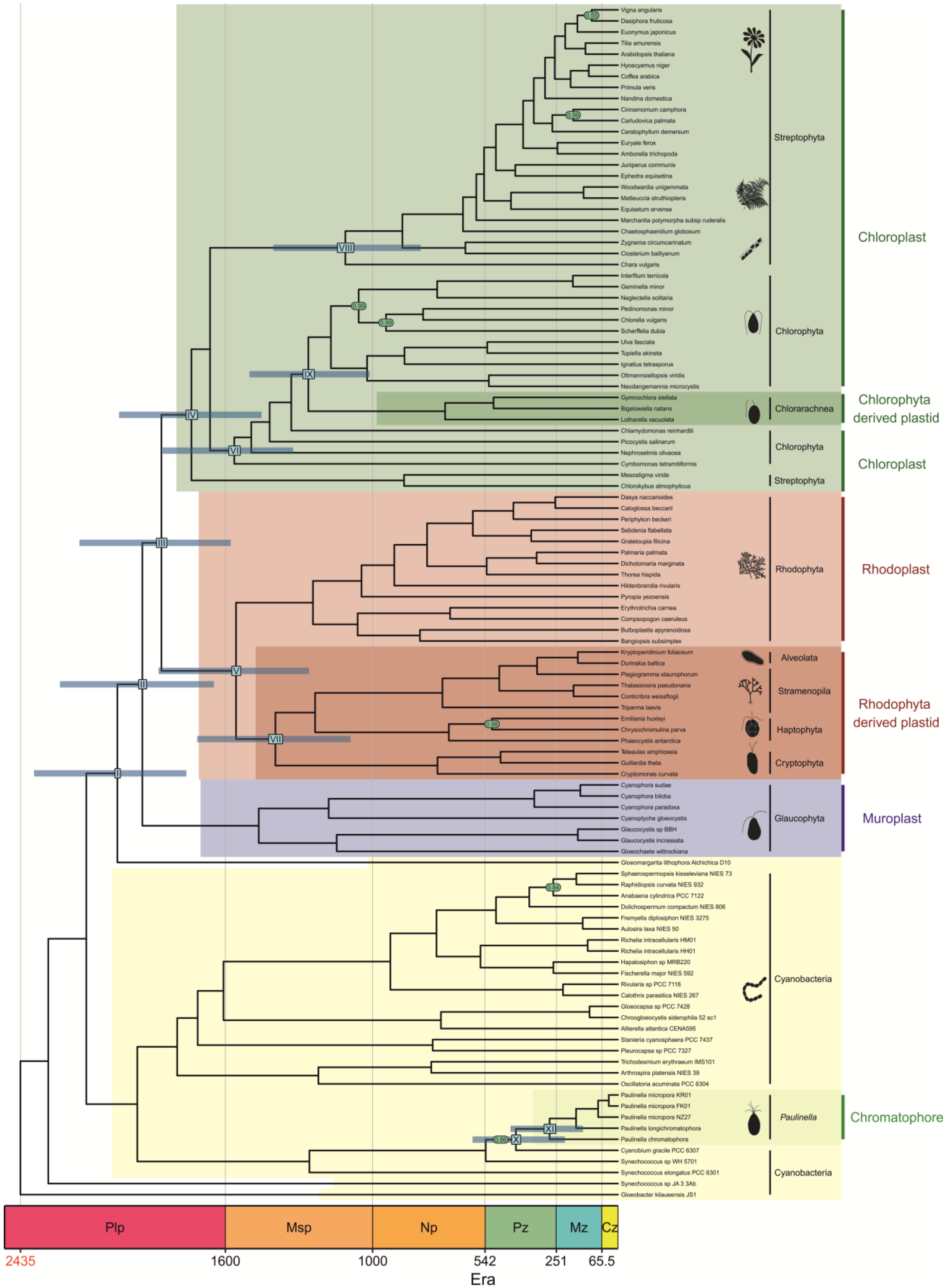


**Figure S13.** Time-calibrated phylogeny of photosynthetic organelles and cyanobacteria. The tree was inferred with MrBayes under IGR model and calibrated with C2 (Table 1). Other description as in Fig. S1.

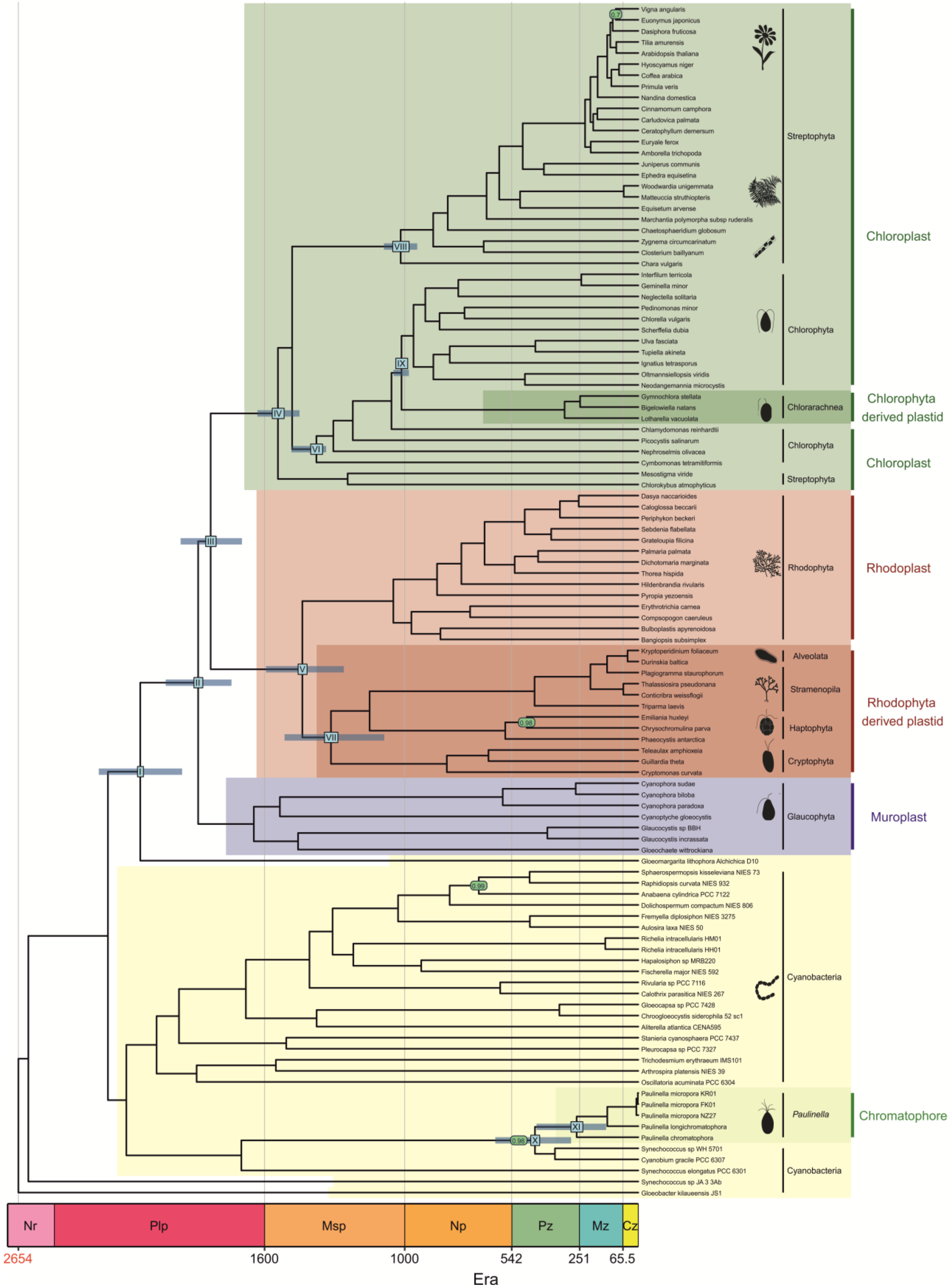


**Figure S14.** Time-calibrated phylogeny of photosynthetic organelles and cyanobacteria. The tree was inferred with MrBayes under TK02 model and calibrated with C2 (Table 1). Other description as in Fig. S1.

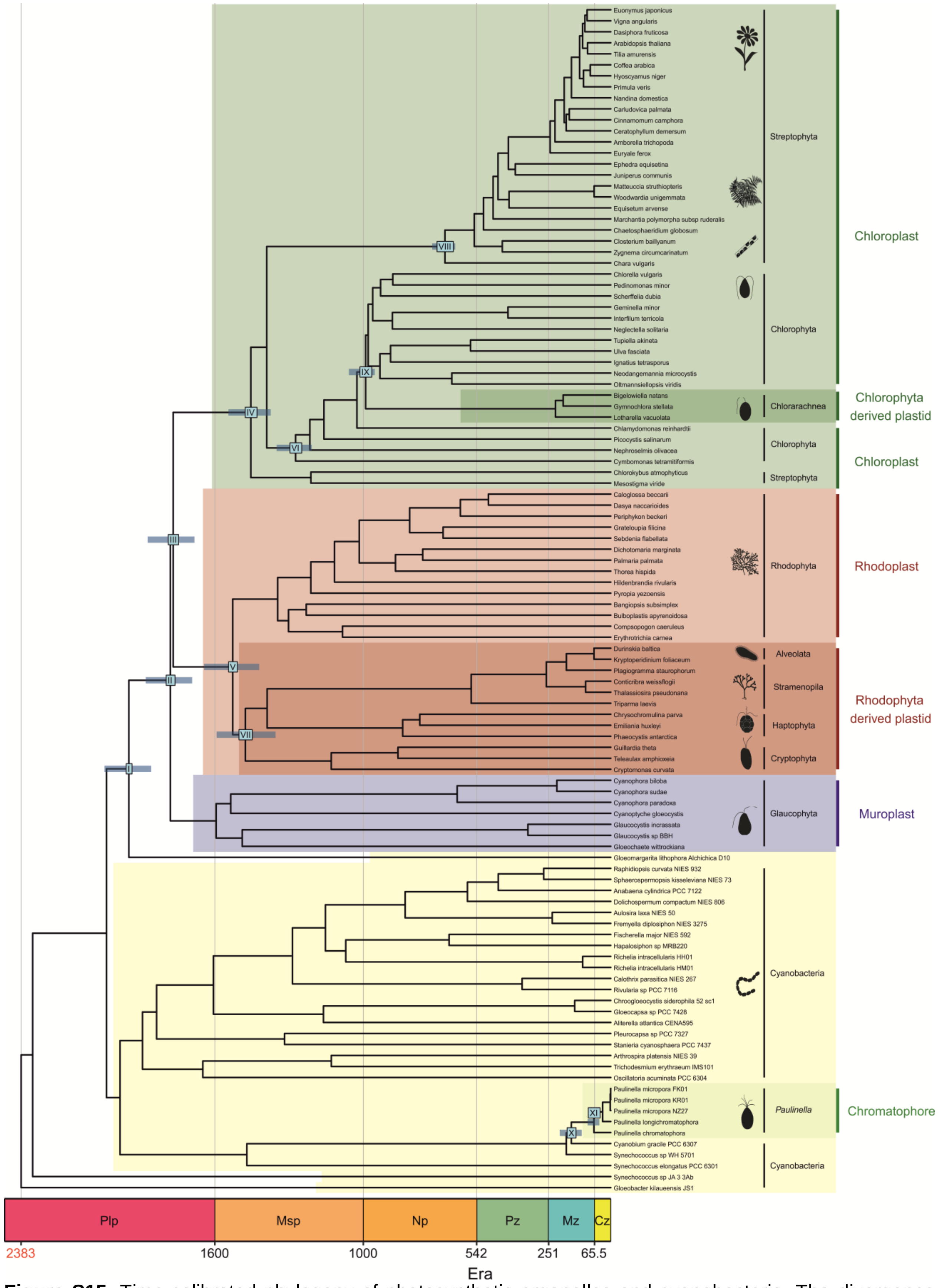


**Figure S15.** Time-calibrated phylogeny of photosynthetic organelles and cyanobacteria. The divergence times were inferred with Phylobayes under CIR model and calibrated with C2 (Table 1). The tree topology was reconstructed in IQ-TREE. Other description as in Fig. S1.

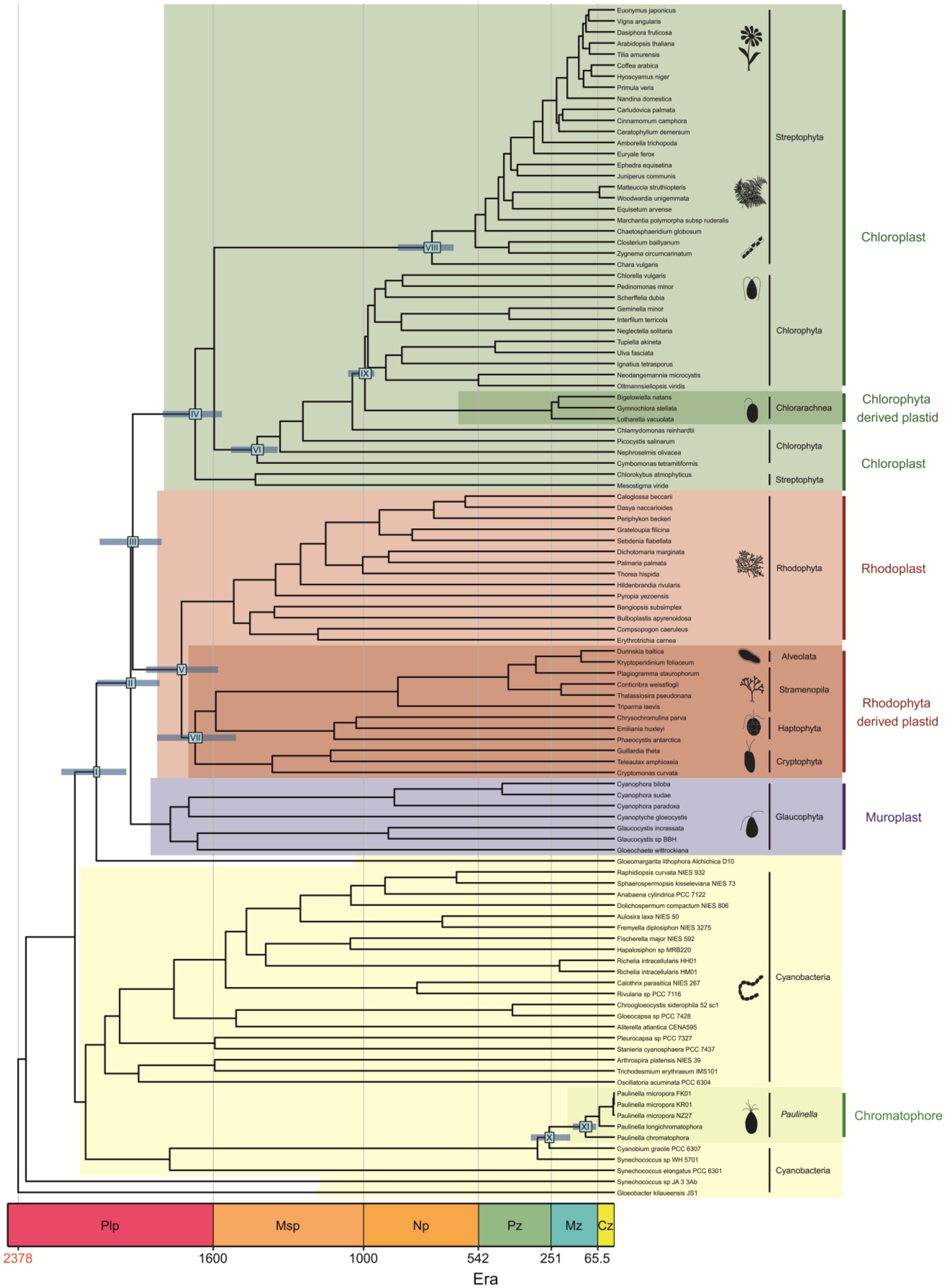


**Figure S16.** Time-calibrated phylogeny of photosynthetic organelles and cyanobacteria. The divergence times were inferred with Phylobayes under LN model and calibrated with C2 (Table 1). The tree topology was reconstructed in IQ-TREE. Other description as in Fig. S1.

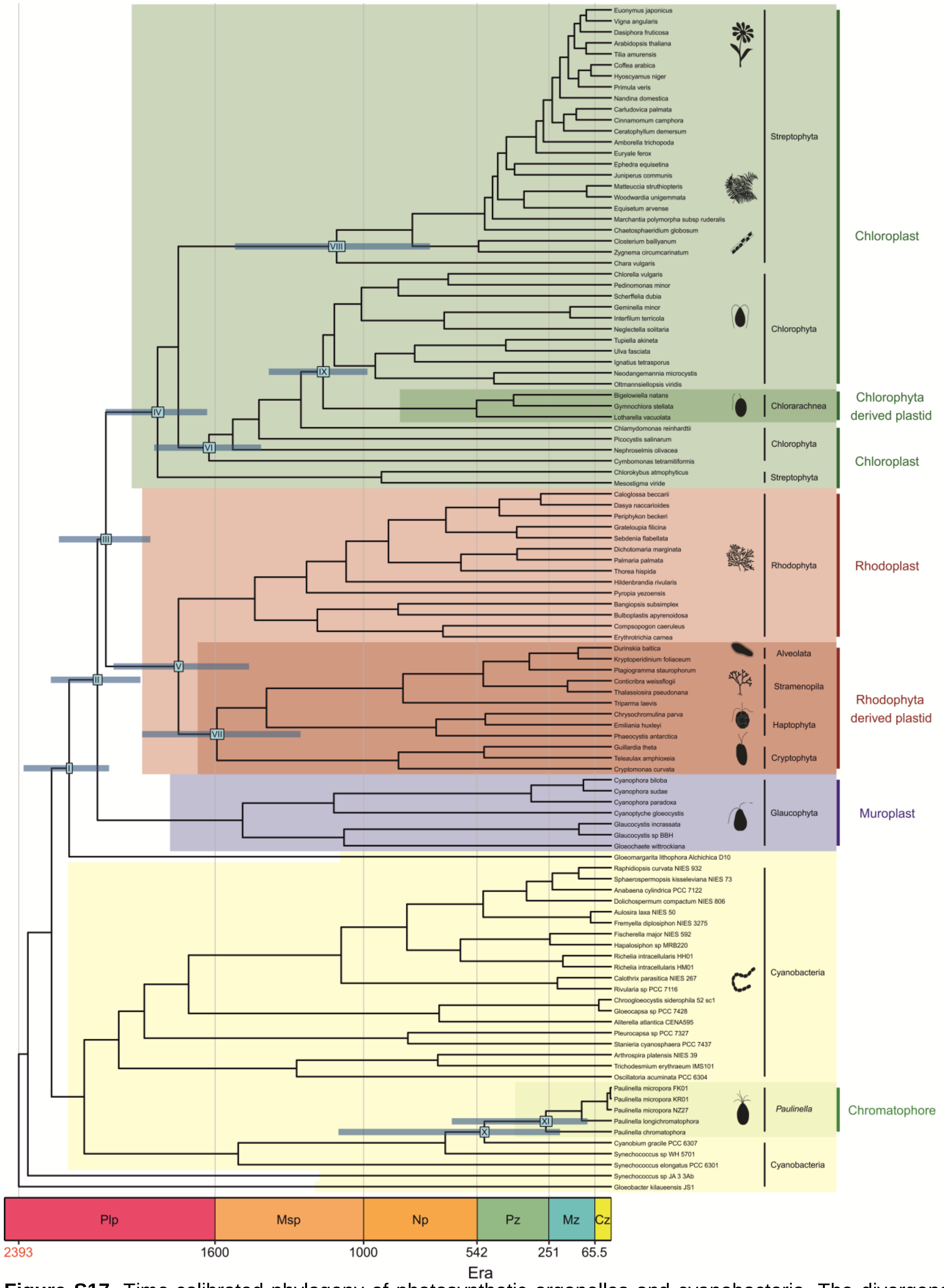


**Figure S17.** Time-calibrated phylogeny of photosynthetic organelles and cyanobacteria. The divergence times were inferred with Phylobayes under UGAM model and calibrated with C2 (Table 1). The tree topology was reconstructed in IQ-TREE. Other description as in Fig. S1.

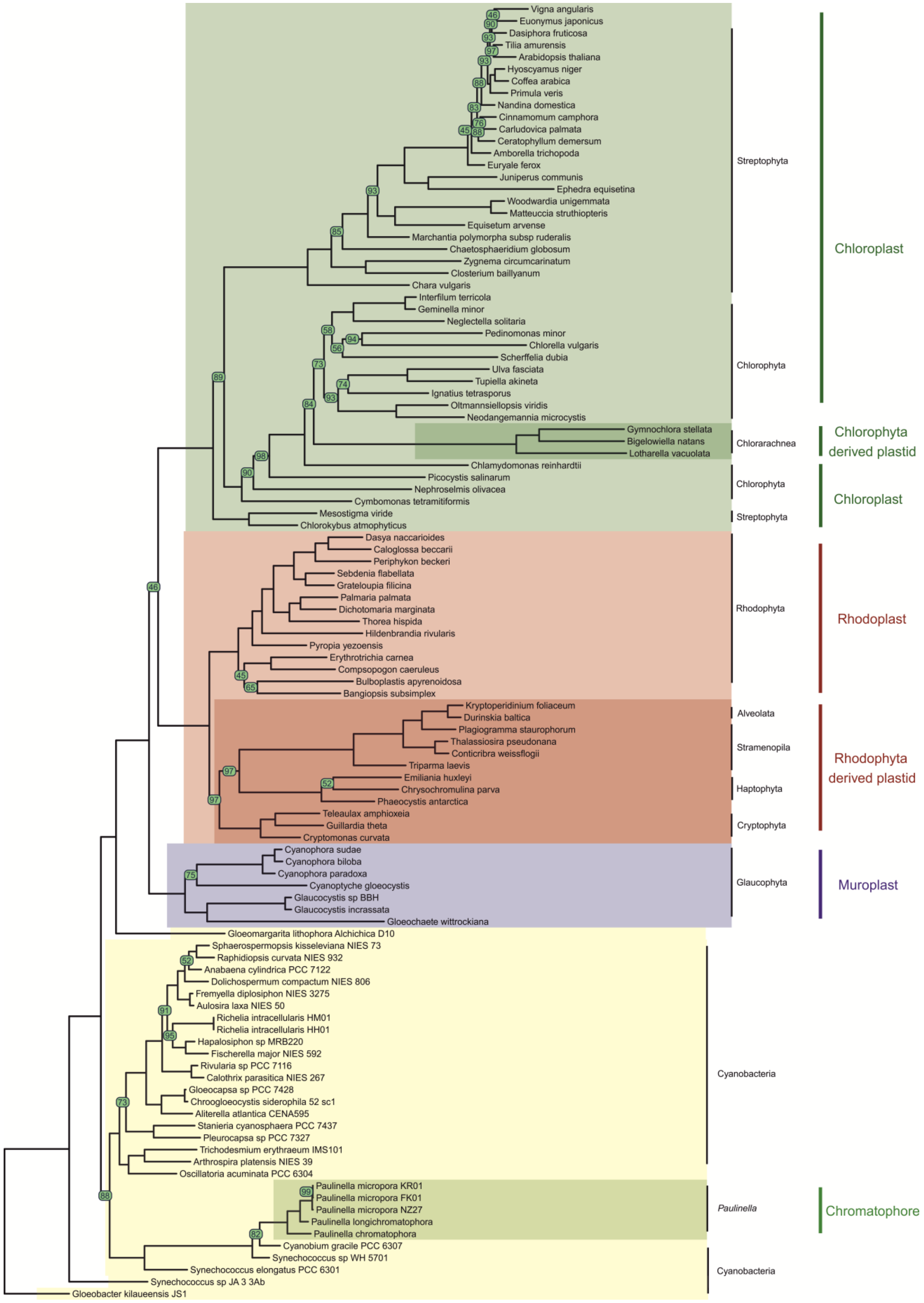


**Figure S18.** The phylogeny of photosynthetic organelles and cyanobacteria. The tree topology was reconstructed in IQ-TREE. The nodes supported with bootstrap values lower than 100 are indicated in green circles.

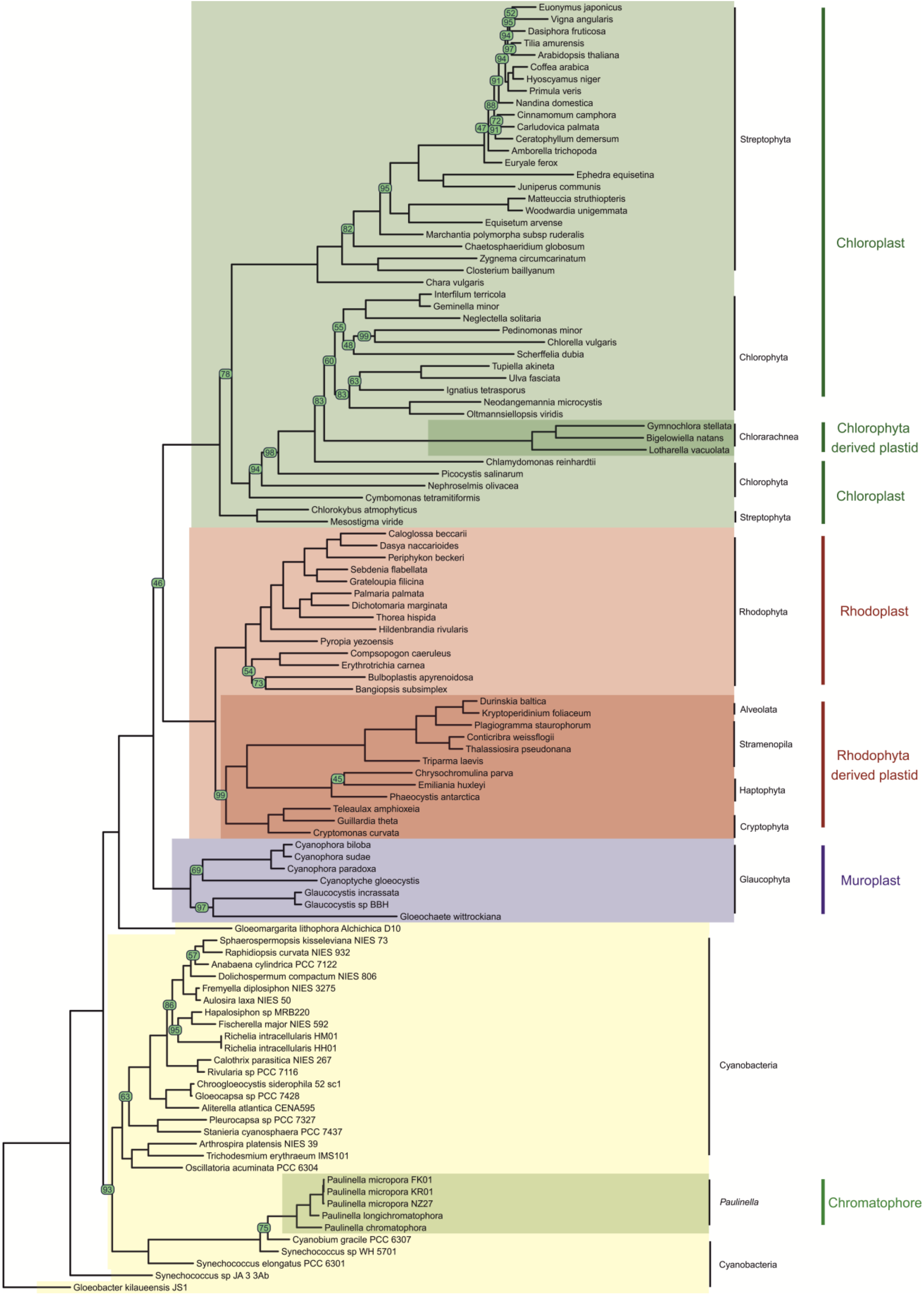


**Figure S19.** The phylogeny of photosynthetic organelles and cyanobacteria. The tree topology was reconstructed in RAxML. The nodes supported with bootstrap values lower than 100 are indicated in green circles.

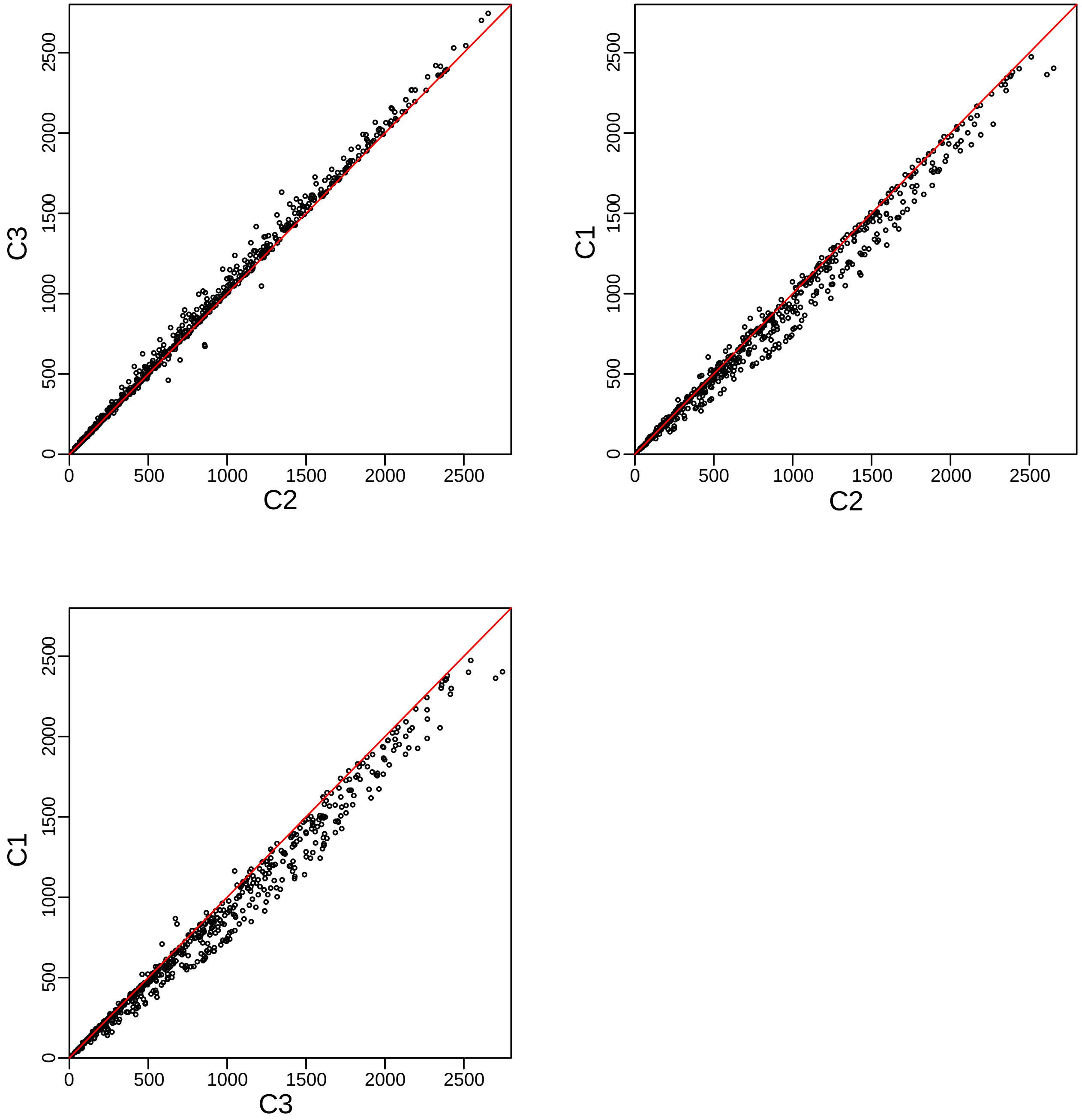


**Figure S20.** Pairwise comparisons of the calibration sets C1, C2 and C3 (Tab.1). The x and y-axis represent ages in million years. The circles are individual age estimations for a node with a given molecular clock and calibration set.

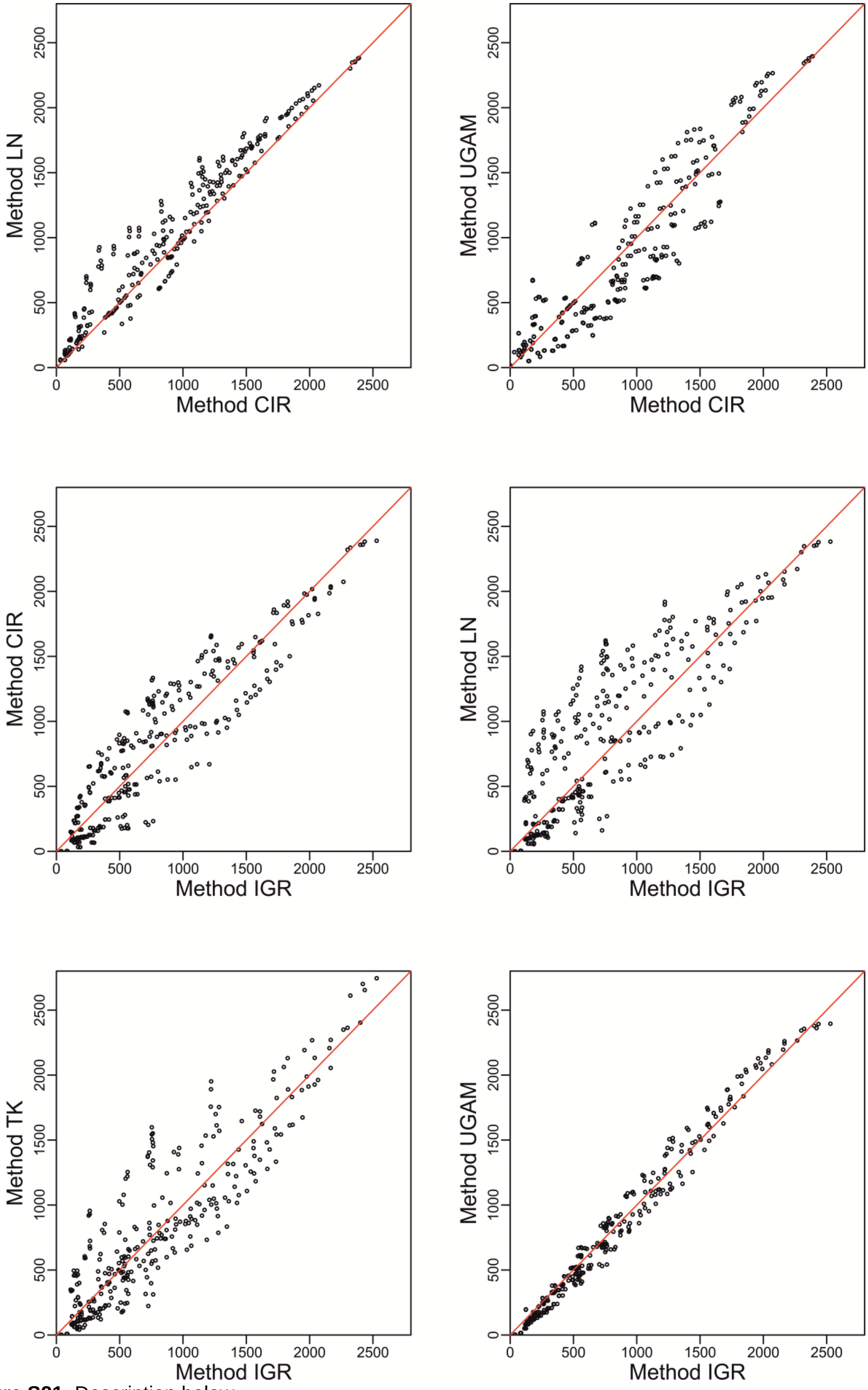

**Figure S21.** Description below.

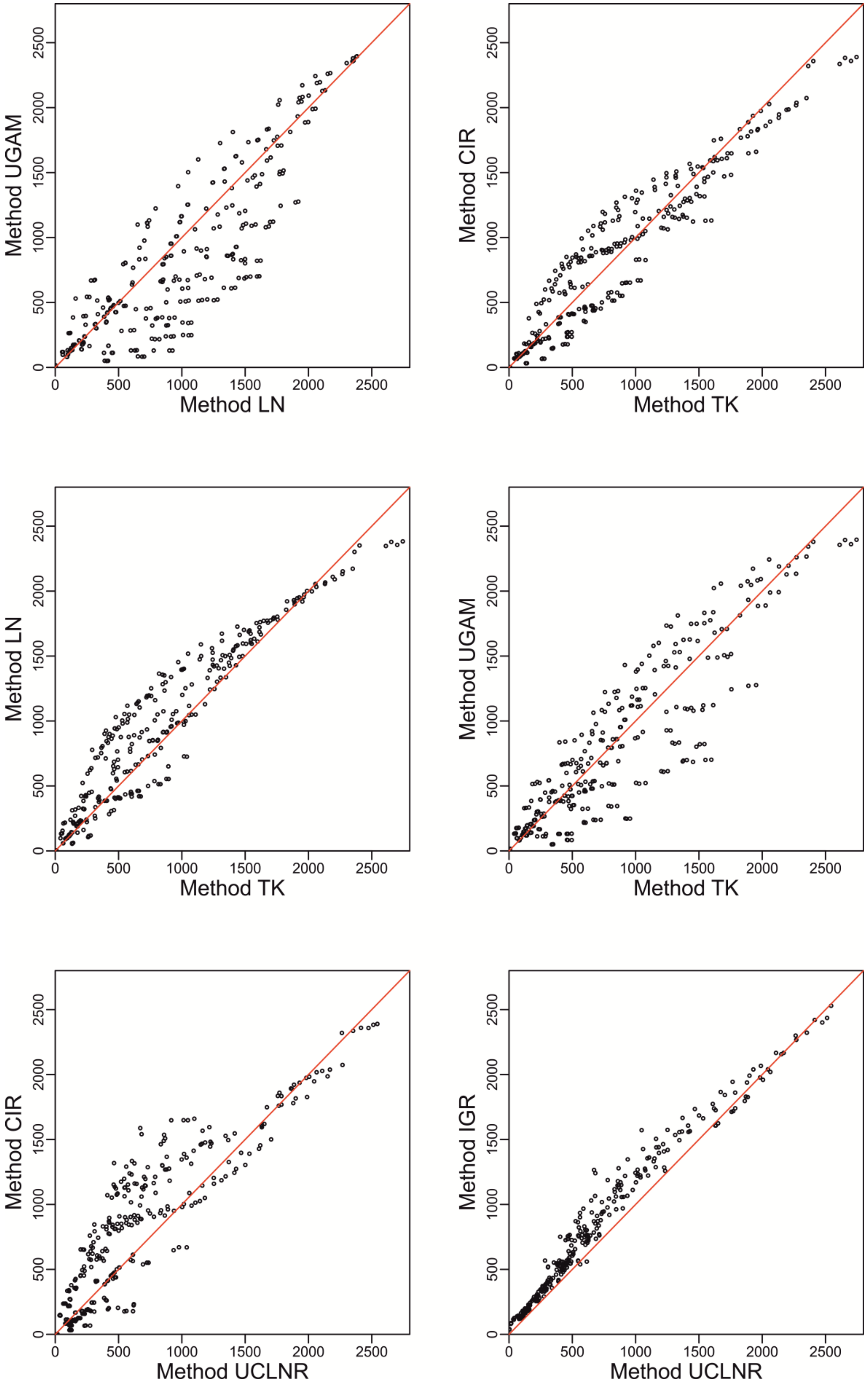

**Figure S21.** Description below.

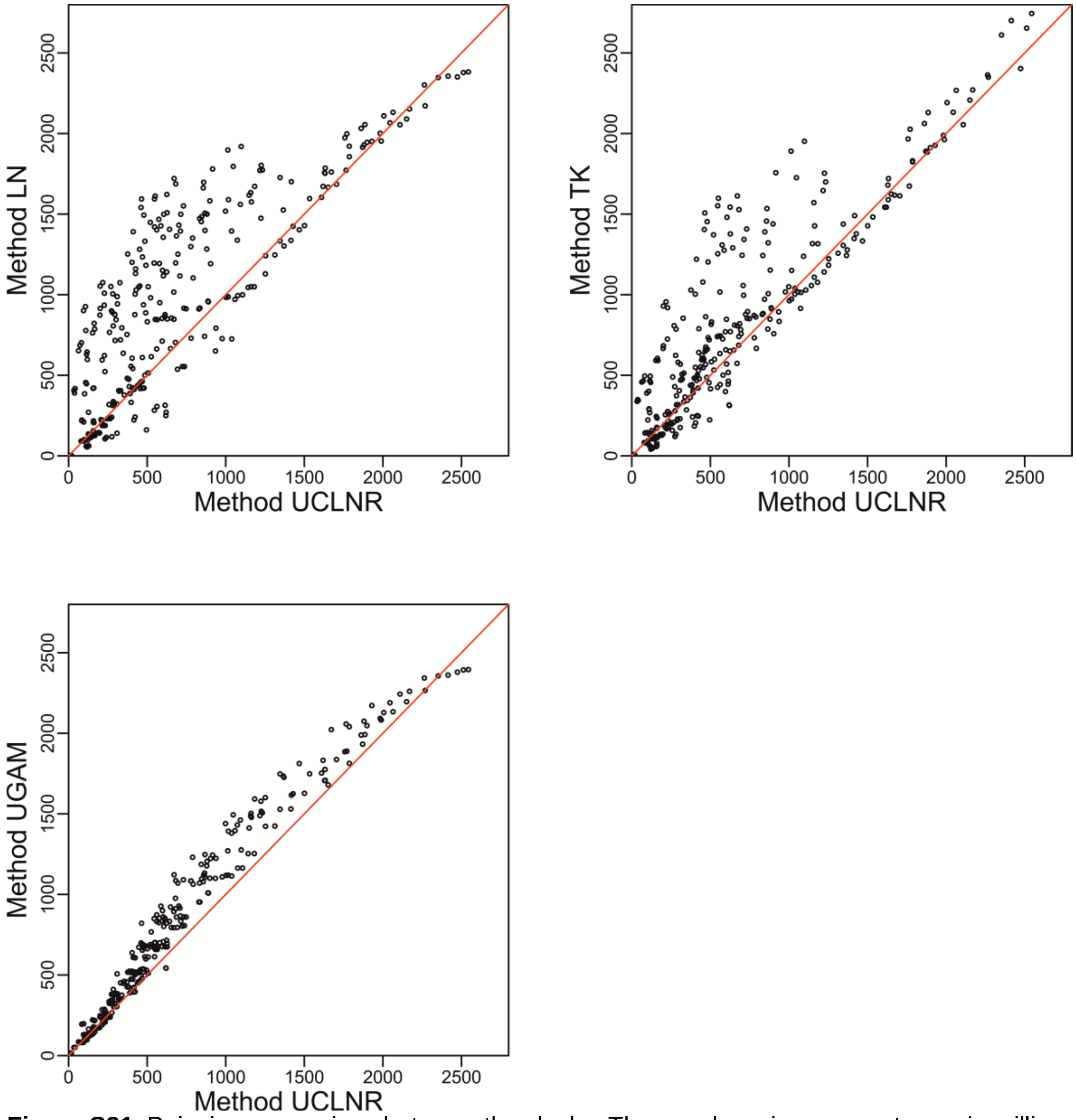


**Figure S21.** Pairwise comparison between the clocks. The x and y-axis represent ages in million years. The circles are individual age estimations for a node with a given molecular clock and calibration set.
